# RNAi screening identifies a new Toll from shrimp that restricts WSSV infection through activating Dorsal to induce antimicrobial peptides

**DOI:** 10.1101/365197

**Authors:** Haoyang Li, Bin Yin, Sheng Wang, Qihui Fu, Bang Xiao, Kai Lǚ, Jianguo He, Chaozheng Li

## Abstract

The function of Toll pathway defense against bacterial infection has been well established in shrimp, however how this pathway responds to viral infection is still largely unknown. In this study, we report the Toll4-Dorsal-AMPs cascade restricts the white spot syndrome virus (WSSV) infection of shrimp. A total of nine Tolls from *Litopenaeus vannamei* namely Toll1-9 are identified, and RNAi screening *in vivo* reveals the Toll4 is important for shrimp to oppose WSSV infection. Knockdown of Toll4 results in elevated viral loads and renders shrimp more susceptible to WSSV. Furthermore, Toll4 could be a one of upstream pattern recognition receptor (PRR) to detect WSSV, and thereby leading to nuclear translocation and phosphorylation of Dorsal, the known NF-κB transcription factor of the canonical Toll pathway. More importantly, silencing of Toll4 and Dorsal contributes to impaired expression of a specific set of antimicrobial peptides (AMPs) such as anti-LPS-factor (ALF) and lysozyme (LYZ) family, which exert potent anti-WSSV activity. Two AMPs of ALF1 and LYZ1 as representatives are demonstrated to have the ability to interact with several WSSV structural proteins. Taken together, we therefore identify the Toll4-Dorsal pathway mediates strong resistance to WSSV infection by inducing some specific AMPs.

**Author summary:** The TLR pathway mediated antiviral immune response is well identified in mammals, yet, Toll pathway governing this protection in invertebrates remains unknown. In the present study, we uncover that a shrimp Toll4 from a total of nine Tolls in *L. vannamei* confers resistance to WSSV thought inducing the NF-κB transcription factor Dorsal to inspiring the production of some antimicrobial peptides (AMPs) with antiviral activity. The anti-LPS-factor (ALF) and lysozyme (LYZ) family are identified as the Toll4-Dorsal pathway targeted genes with the ability to interact with viral structural proteins in response to WSSV infection. These results suggest that the Toll receptor induces the expression of AMPs with antiviral activity could be a general antiviral mechanism in invertebrates and Toll pathway established antiviral defense could be conserved during evolution.

## Introduction

Multicellular organisms have evolved the ability to protect themselves from a wide variety of pathogens including virus. In invertebrates including shrimps that lacking immunoglobulin-based adaptive immune system, this protection is thus provided through the action of an innate immune system. The innate immune response is generally initiated via the detection of pathogen-associated molecular patterns (PAMPs), some evolutionarily conserved structures or motifs shared by broad classes of invading organisms, by a wide diversity of host pattern recognition receptors (PRRs) (1). One important class of PRRs is the Toll receptor superfamily, comprising invertebrate Tolls and vertebrate Toll-like receptors (TLRs), and is now considered to be the primary sensor of pathogens in all metazoans (2).

Mammalian TLRs play universal and pivotal roles in host defenses mainly via the innate immune system, but also the immunoglobulin-based adaptive immune system that are devoid in invertebrate (3). In human, the function of TLR signaling pathway is clearly clarified, and the ten TLRs can directly recognize and bind to a number of diverse molecular structures, including lipids (e.g., TLR4: LPS via MD2; TLR1/2/6: lipoproteins), proteins (e.g., TLR5: flagellin; TLR2 and TLR4: HMGB1), and nucleic acids (e.g., TLR3: dsRNA; TLR7/8: ssRNA; and TLR9: unmethylated CpG motifs in bacterial, viral, and fungal DNA) (4). It is generally accepted that TLRs require ligand-induced dimerization or crosslinking to initiate intracellular signaling event (4). Ligand binding is likely to lead to a conformational rearrangement of the cytoplasmic TIR domains, thereby creating a docking site to which TIR domain containing adaptors MyD88/TIRAP/TRIF/TRAM can be recruited (2, 4). The stimulation of TLRs then result in activation of the NF-κB (nuclear factor κB) transcription factors that drive the transcription of proinflammatory cytokines, or/ and trigger IRF (interferon regulatory factor) transcription factors that induce transcription of type I interferon cytokines, both of which ultimately confer immune response against infection (2, 4).

*Drosophila* genome encodes nine Toll receptor genes (Toll1 to Toll9), but these Tolls are different and elusive in the context of function, ligand sensing and intracellular signaling compared to those of mammals. *Drosophila* Toll1 (or simply, Toll) generally binds an endogenous cytokine Spätzle (Spz) rather than microbe motif (5). In contrast to that mammalian TLRs function mainly in immunity, *Drosophila* Toll1 functions not only in developmental processes (6, 7), but also in innate immunity response to bacterial, fungal, and viral infections (8-10). *Drosophila* Toll7 can directly interact with vesicular stomatitis virus (VSV) virion at the plasma membrane perhaps in a manner similar to mammalian TLRs, and induces antiviral autophagy independent of the canonical Toll signaling pathway (MyD88/Tube/Pelle/Dorsal or Dif cascade) (11). Besides, *Drosophila* Toll2 (18-wheeler) may play a minor role in the antibacterial response (12), and Toll5 and Toll9 can trigger the production of the antifungal gene Drosomycin (13). By now, only the *Drosophila* Toll1 is definitely identified as upstream receptor of Dorsal or Dif, two transcription factors homologous to *Human* NF-κB (14). However, the function and intracellular signaling routes of the remaining Tolls in response to infection are not well characterized.

Cultured shrimps frequently suffer from many DNA and RNA viruses, among which white spot syndrome virus (WSSV) has been considered as the most serious threat to shrimp aquaculture industries and caused serious economic losses every year (15, 16). WSSV is a large and enveloped dsDNA virus, which is highly pathogenic and especially virulent on penaeid shrimp (15, 17). There is growing interest in research of the interplay between WSSV and many aspects of shrimp host (18, 19), but the precise function of Toll receptors and Toll pathway related genes participating in viral infection remains to be determined. Until now, a large number of Tolls have been identified in multiple shrimps including *Litopenaeus vannamei* (20, 21), *Fenneropenaeus chinensis* (22), *Penaeus monodon* (23-25), *Marsupenaeus japonicas* (26), *Macrobrachium rosenbergi* (27), *Cherax quadricarinatus* (28) and *Procambarus clarkii* (29, 30). In *L. vannamei*, three Tolls of the Toll1, Toll2 and Toll3 are up-regulated after WSSV challenge, whereas their functions during WSSV infection are not well characterized (21). Some shrimp Tolls from *C. quadricarinatus, P. clarkii* and *M. rosenbergii* have been shown to induce some AMPs expression in response to WSSV infection, which indicate these AMPs could play an antiviral role (28, 29, 31). Besides, WSSV infection can contribute to activation of many *P. monodon* Toll pathway genes, which suggest that the whole pathway play a crucial role in the immune response during WSSV infection (32). In a recent study, the Toll3 from *L. vannamei* is demonstrated to have the ability to activate the expression of interferon regulatory factor (IRF) and its downstream Vago4/5, suggesting it could play a critical role in host antiviral immunity independent of the canonical Toll signaling pathway (33). Overall, some shrimp Tolls have been shown to participate in innate immune responses against viral infection, but their antiviral functions remain largely elusive and their underlying antiviral mechanism needs further investigation clearly.

Herein, we clone and identify a total of nine Tolls (Toll1-9) from *L. vannamei*, and RNAi screening demonstrates Toll4 as a critical antiviral factor against WSSV *in vivo*. Mechanismly, Toll4 senses WSSV infection and leads to activation of Dorsal to converge on the production of some specific AMPs such as ALFs and LYZs, which confer host defense against WSSV by targeting its structural proteins. These data provide evidence that a new identified Toll4 senses WSSV and initiates an antiviral response in shrimp.

## Results

### Cloning, sequence analysis and phylogenetic tree of shrimp nine Tolls

In order to identify all candidate Toll genes from shrimp *(L. vannamei)*, we carried out protein homology search by local TBLASTN program against our transcriptome data from whole bodies of *L. vannamei* (all tissues pooled) (34) and other *L. vannamei* transcriptome data from NCBI (35-38). A total of nine individual and putative Toll homologs from *L. vannamei* were obtained, of which the Toll1, Toll2 and Toll3 were perfectly identical to previous reported LvToll1, LvToll2 and LvToll3, respectively (21). We next cloned the full-length cDNA sequences of other Tolls by using rapid amplification cDNA ends (RACE)-PCR method, and we subsequently designated these Tolls from Toll1 to Toll9. Their sequences of Toll1-9 in FASTA format were available in Supplement Data S1. Functional domain analysis indicated that each of the nine Tolls adopted a typical domain organization characteristic of Toll family gene including an N-terminal signal peptide, an extracellular domain, a single transmembrane region and an intracellular TIR domain in the C-terminal (Figure 1A). These Tolls varied considerably in the number of extracellular LRRs from 7 (Toll7) to 28 (Toll5) (Figure 1A), and the intracellular TIR domains were not conserved among each other except for the two pairs of Toll1/2 and Toll3/Toll8 bearing sequence identities of greater than or equal to 70% (Figure S1). The significantly different both in number of extracellular LRRs and sequence identities of intracellular TIR domains may suggest that these Tolls are able to respond to various extracellular stresses (pathogens or ligands) and exploit distinct intracellular signaling routes. Phylogenetic tree analysis showed that these Toll and TLR homologs were divided into three clusters: the invertebrate cluster including most of Tolls from shrimps and other invertebrates, the vertebrate cluster including TLRs from mammals and the *Drosophila* Toll9 as well as *L. vannamei* Toll9, and the another cluster consisting of a sole *L. vannamei* Toll7 (Figure 1B). Taken together, we cloned and identified nine Tolls from *L. vannamei*, among which six Tolls namely the Toll4, Toll5, Toll6, Toll7, Toll8 and Toll9 are firstly cloned and identified in *L. vannamei*.

**Figure 1.**
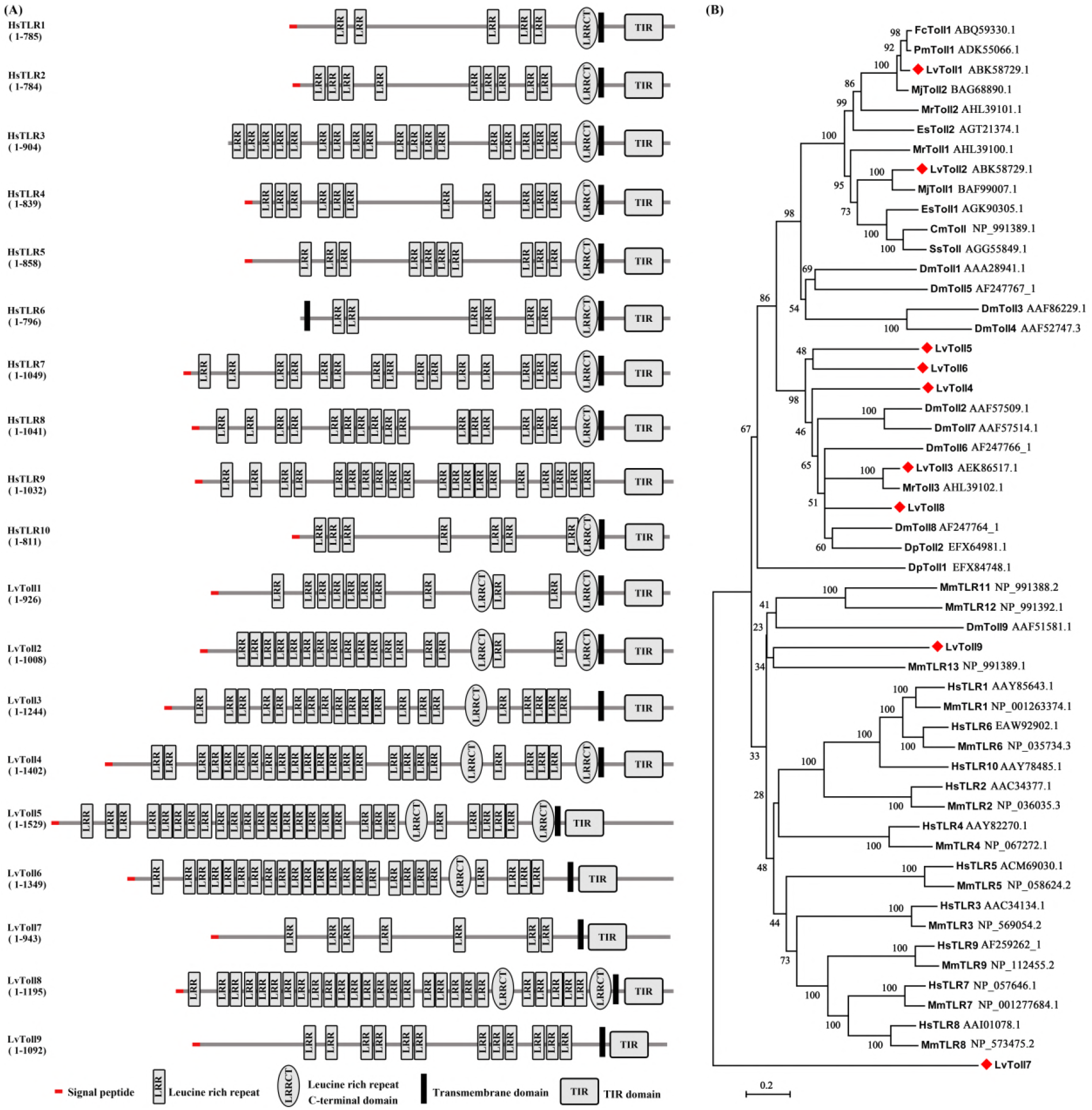
Structural organization and phylogenetic analysis of shrimp Toll1-9 and Tolls/Toll-like receptors (Tolls/TLRs) from other species. (A) Schematic representations of the domain topology of *L. vannamei* Tolls (LvToll1–9) and human TLRs (HsTLR1–10) according to SMART analysis. (B) Phylogenetic tree of Tolls/TLRs. The tree was constructed with the neighbour-joining (NJ) method based on the alignment of 53 Toll/TLRs full-length protein sequences by utilizing MEGA 5.0 software. The bootstrap values of 1000 replicates (%) were indicated on the branch nodes. *L. vannamei* nine Tolls (LvToll1-9) were indicated in red diamonds. More detail information of sequences about these Tolls/TLRs was supplied with the Supplement Data S3.

### Toll4 and the canonical Toll pathway components restrict WSSV infection in shrimp

To determine whether any of the shrimp Tolls are involved in antiviral defense against WSSV, we generated double-stranded RNA (dsRNA) against each of the nine Tolls and silenced them *in vivo* via dsRNA mediated RNA interference (RNAi). We firstly addressed the tissue distribution of shrimp nine Tolls transcripts by Semi-quantitative reverse transcription PCR (Semi-qRT-PCR). The results showed that both Toll1 and Toll4 could be detected in all the examined tissues and were highly expressed in gill, hemocyte and intestine, whereas other Tolls were abundant in only a few specific tissues (Figure 2A). According to the tissue distribution of each Tolls, the gill tissue was chosen to check the knockdown efficiencies for Toll4, Toll6, Toll7 and Toll8 (Figure 2C), while hemocyte was as the target tissue to evaluate silencing efficiencies for Toll1, Toll2, Toll3 Toll5 and Toll9 (Figure 2B). Efficient silencing for each Toll receptor was confirmed by quantitative reverse transcription PCR (qRT-PCR) (Figures 2B and 2C). Next, we challenged RNAi-treated shrimps with WSSV and subsequently analyzed viral genome copies (WSSV DNA) by absolute quantitative PCR (absolute q-PCR). We observed a greater number of each Toll expect for Toll2 silenced shrimps exhibited higher quantities of viral titers in muscle when compared to control shrimps at 48 hours post infection (hpi) (Figure 2D). Of note, shrimps with silencing of Toll4 had the highest WSSV titers, and the median of viral DNA burden was approximately 150 times higher than that of the control shrimps (GFP dsRNA injected shrimps following WSSV infection) (Figure 2D). Therefore, we focused our attention special on the Toll4 mediated mechanism underlying resistance to WSSV in this study. To further confirm the above screening result, with *in vivo* RNAi again, a higher lethality was observed in the silencing of Toll4 shrimps followed by WSSV infection when compared to that of the control group (Figure 2E). To investigate whether the increased lethality rates of Toll4 silenced shrimps was due to decreased resistance or tolerance to WSSV, we analyzed the viral levels by absolute q-PCR in several tissues including hepatopancreas, gill and muscle at 48 hpi. We observed that shrimps with knockdown of Toll4 had elevated viral replication levels in all the three tissues than control shrimps (Figure 2F). The obviously lower viral levels observed in hepatopancreas than those of gill and muscle can be explained by that only the connective tissue and myoepithelial cells of the hepatopancreas sheath are infected by WSSV (39, 40). Collectively, these data strongly suggest Toll4 as a critical antiviral factor against WSSV *in vivo*.

**Figure 2.**
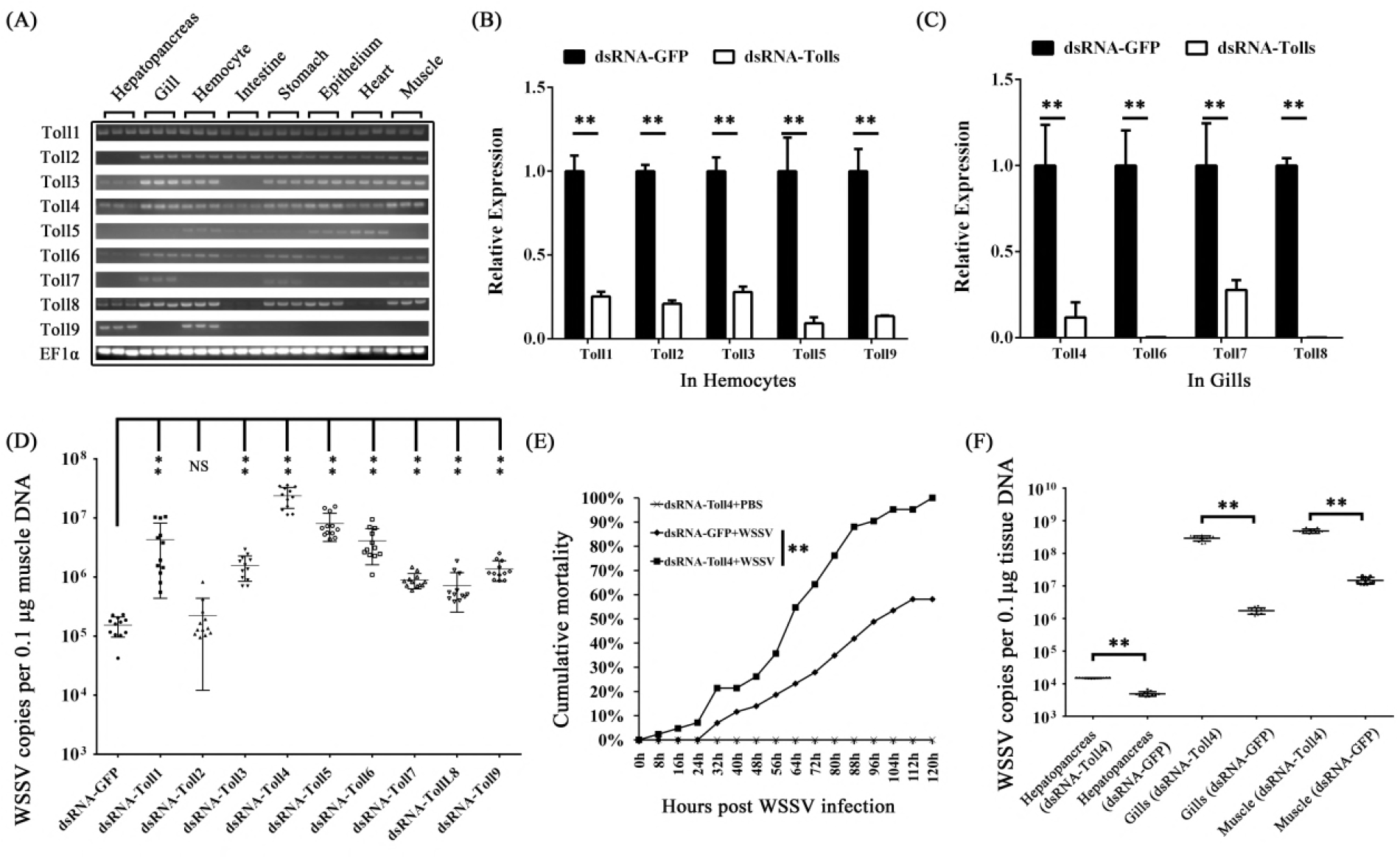
RNAi screening identifies Toll4 as an antiviral factor against WSSV. (A) Tissue distribution of *L. vannamei* Tolls (Toll1-9) was analyzed by Semi-RT-PCR, and EF1α was used as a control. (B-C) Knockdown efficiencies of Toll1, Toll2, Toll3, Toll5 and Toll9 in hemocytes (B), and silencing efficiencies of Toll4, Toll6, Toll7 and Toll8 in gills (B) were checked by qRT-PCR. The GFP dsRNA treated shrimp was set as a control. (D) RNAi screening identified shrimp Tolls (LvToll1-9) as potential antiviral (anti-WSSV) factors. WSSV was inoculated at 48 h post each Toll silencing. The viral load was assessed at 48 h post-infection through absolute qPCR. The experiment was repeated three times with similar results. One dot represented 1 shrimp and the horizontal line represented the median of the results. (E) Silencing of Toll4 enhanced shrimps susceptibility to WSSV infection. WSSV was inoculated at 48 h post Toll4 silencing, and the death of shrimp was recorded at every 8 h for cumulative mortality rates analysis. The experiment was repeated three times, and similar results were obtained. The data were analyzed statistically by the Kaplan–Meier plot (log-rank χ^2^ test) (\*\**p* < 0.01). (F) Silencing of Toll4 enhanced WSSV replication in multiply shrimp tissues. WSSV was inoculated at 48 h post Toll4 silencing, and the viral load was assessed at 48 hpi through absolute qPCR. The experiment was performed three times with similar results. All the data from B, C, D and F were analyzed statistically by student’s T test (\*\**p* < 0.01; NS, not significant).

Because the Toll4 can confer protection against WSSV, we then explored whether the canonical Toll pathway components including MyD88, Tube, Pelle and Dorsal are involved in antiviral responses. We injected shrimps with dsRNA of each component to depress their expression, which were confirmed by quantitative RT-PCR (Figure 3A). We observed that knockdown of these critical Toll pathway components had notable impact on WSSV replication *in vivo* (Figure 3B), suggesting that the canonical Toll pathway plays an important role against WSSV. In summary, these data showed that Toll4 and the canonical Toll pathway components are important to oppose WSSV infection.

**Figure 3.**
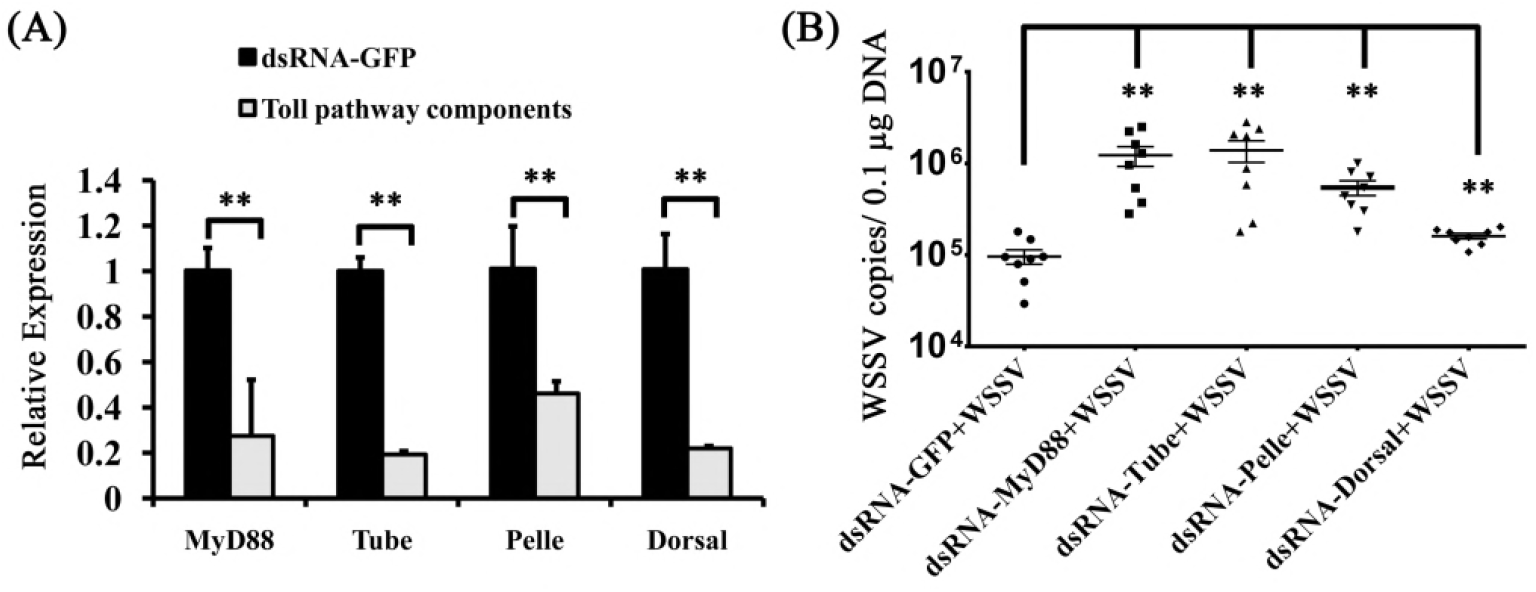
Increased viral replication levels in the canonical Toll pathway components silenced shrimps. (A) Knockdown efficiencies of the canonical Toll pathway components including MyD88, Tube, Pelle and Dorsal in gills were checked by qRT-PCR. The GFP dsRNA treated shrimp was set as a control. (B) Silencing of the canonical Toll pathway components resulted in enhanced WSSV replication levels in gill tissues. WSSV was inoculated at 48 h post each component silencing, and the viral load was assessed at 48 hpi through absolute qPCR. The experiment was performed three times with similar results. All the data were analyzed statistically by student’s T test (** *p* < 0.01).

### Toll4 regulates antimicrobial peptides (AMPs) production upon WSSV infection in shrimp

Mammalian TLRs opposed viral infection mainly through inducing the type I interferon (IFN) expression (41), whereas *Drosophila* Toll restricted viral infection via inspiring some specific AMPs production (10). To address whether the involvement of Toll4 in regulating AMPs synthesis, we firstly studied the response of Toll4 to viral infection and measured the transcriptional changes of Toll4 after WSSV challenge in the two immune related tissues gill and hemocyte by quantitative RT-PCR. The results showed that Toll4 in gill was remarkably up-regulated from 8 hpi to 24 hpi compared to the control shrimps injected only PBS (Figure 4A), and Toll4 in hemocytes maintained an elevated expression level in the whole period of infection (Figure 4B). To test whether some known shrimp AMPs can respond to WSSV infection, we detected the transcriptional levels of fourteen shrimp AMPs consisting of four different AMP families of anti-lipopolysaccharide (LPS) factor (ALF), lysozyme (LYZ), penaeidin (PEN) and crustin (CRU), by quantitative RT-PCR at 6 hpi in gill and hemocyte tissues (Figures 4C and 4D). These sequences of fourteen shrimp AMPs in FASTA format were available in Supplement Data S2. We used *Vibrio parahaemolyticus*, a bacterial pathogen of shrimp that has been verified as an activator of the shrimp canonical Toll-Dorsal pathway (42), as a control to compare the levels of AMPs expression in viral infection to those in this bacterial infection. PBS controls were used for baseline levels of AMP expression. We found that, except for ALF4, PEN2 and PEN3 with slight a up-regulation less than 2-fold in gills, both WSSV and *V. parahaemolyticus* infection induced considerably increased expression levels of the most of AMPs over 2-fold than those of control shrimps in the both tissues (Figures 4 and 4D). Moreover, we used the RNAi to knockdown the expression of Toll4 *in vivo* once again, and the silencing efficiencies of Toll4 in gill and hemocyte at 24 h and 48 h post dsRNA injection were further confirmed by quantitative RT-PCR (Figures 4E and 4F). After 48 h post dsRNA injection, we infected the RNAi treated shrimps by injection of WSSV, and probed the expression levels of the fourteen AMPs at 6 hpi. The results showed that all of these AMPs, expect for CRU3 with a slight up-regulation, were dramatically down-regulated compared to the control in hemocytes (Figures 4H). In contrast to most AMPs of LYZ and ALF families, RNAi of Toll4 did not lead to the down-regulated secretion of other AMPs, except for CRU3, in the gills of WSSV infected shrimps (Figures 4G). These results suggest that Toll4 induced AMPs expression may vary in gill and hemocyte tissues under WSSV infection. Even so, all or most of the tested AMPs from LYZ and ALF families in both hemocytes and gills showed an identical down-regulation pattern in Toll4 silenced shrimps under WSSV infection. In summary, we conclude that WSSV infection in shrimp could activate Toll4 expression and perhaps through Toll4 to induce the production of a specific set of AMP genes including LYZ1-4 and ALF1-4 both in gill and hemocyte tissues.

**Figure 4.**
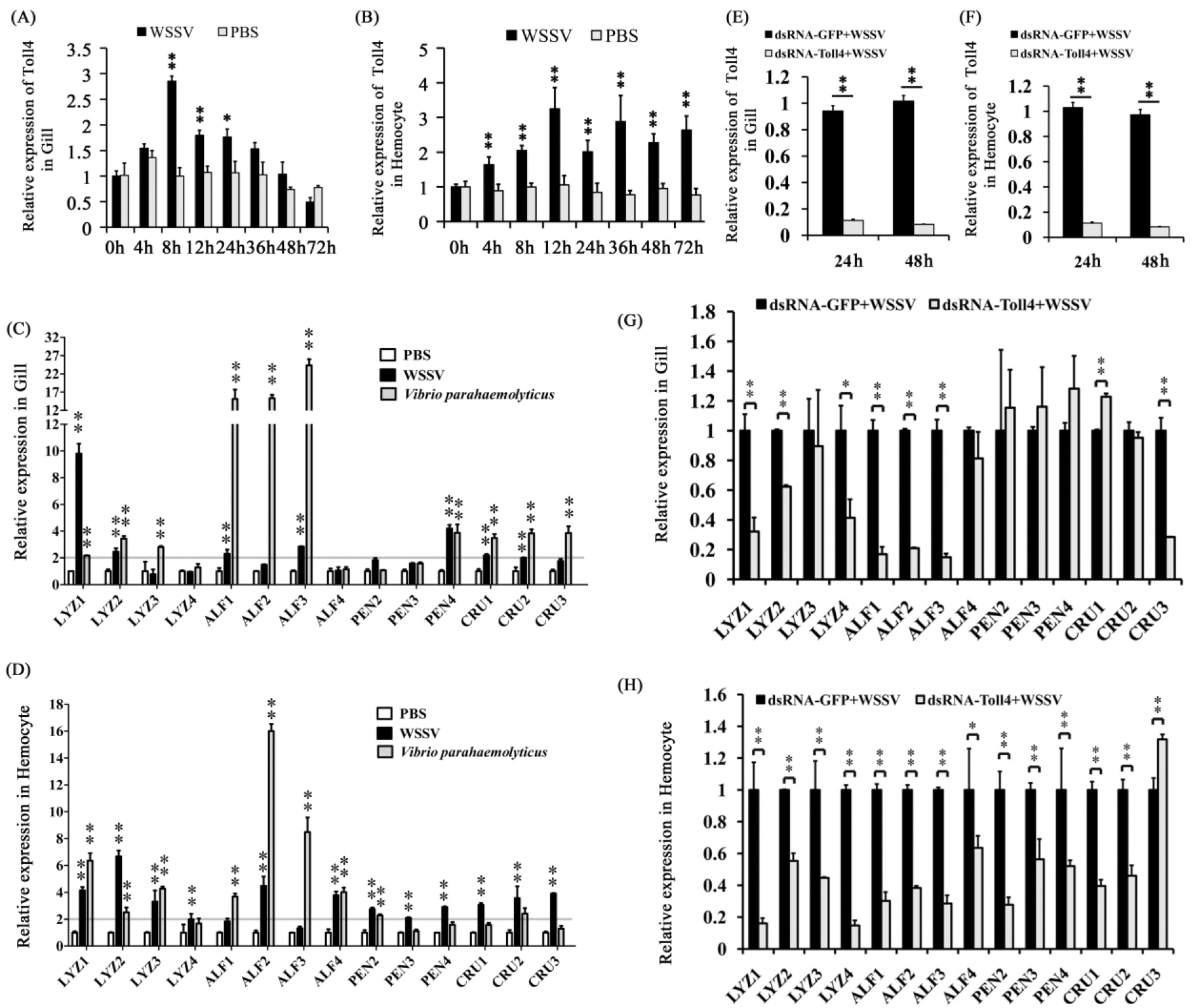
AMPs expression levels in Toll4 silenced shrimps upon WSSV infection. (A-B) Expression profiles of Toll4 after WSSV infection in gill (A) and hemocyte (B) were assessed by qRT-PCR. (C-D) AMPs expression patterns responding to the challenge of WSSV, *V. parahaemolyticus* (as an activator for canonical Toll pathway) and PBS (as a negative control) in gill (C) and hemocyte (D) was detected by qRT-PCR. The horizontal line indicated 2-fold induction threshold. (E-F) Knockdown efficiencies of Toll4 in gill (E) and hemocyte (F) was confirmed by qRT-PCR at 24 and 48 h post dsRNA injection. (G-H) Toll4-knockdown shrimps had impaired AMPs expression levels upon WSSV infection both in gill (G) and hemocyte (H). All experiments were performed three times, and similar results were observed. All the data from A-F were analyzed statistically by student’s T test (* *p* < 0.05; ** *p* < 0.01).

### Dorsal is activated following exposure to WSSV challenge

Since we observed that Toll4 responded to WSSV infection and ultimately led to the induction of a specific set of AMP genes, we explored whether Dorsal, the NF-κB transcription factor known to be downstream of the canonical Toll pathway (43), is activated during viral infection. We firstly detected the tissue distribution of shrimp Dorsal, and found that Dorsal showed high expression levels in gill, hemocyte, stomach, intestine and epithelium, but low in hepatopancreas (Figures 5A). We then investigated the effect of WSSV infection on Dorsal nuclear translocation by immunofluorescence staining in shrimp hemocytes. To gain more information on nuclear translocation of Dorsal upon WSSV infection, we firstly detected the dynamics of Dorsal translocation at several times, and we observed that shrimp hemocytes exhibited gradually and significantly increased nuclear translocation levels of Dorsal (53%, 88% and 94%) compared to the control hemocytes treated with PBS (Normal 0 h, 20%) at 1 h, 3 h and 6 h after WSSV challenge, respectively (Figures 5B and b). To further confirm the above results, we probed Dorsal translocation from the cytoplasm to the nucleus upon WSSV infection by using an *L. vannamei* Dorsal specific antibody prepared in our previous study (44). In good agreement with the results of immunofluorescence staining, we were able to detect more nuclear import of Dorsal in shrimp hemocytes along with the times of WSSV infection from 0 h to 6 h, and at 6 hpi almost no signal of Dorsal can be detected in the cytoplasm (Figure 5C). In addition to nuclear translocation, the activation of Dorsal could be manifested by phosphorylation on some specific amino acids. We found that shrimp Dorsal contains a considerable conserved region, which displays a significant degree of sequence similarity to a comparison region of its mammalian counterparts. In this region, human p65 NF-κB factor contains a Ser276, corresponding to Ser342 of shrimp Dorsal, which can be phosphorylated upon many stresses including viral infection (45) (Figure 5D). We therefore hypothesize that the Ser342 of Dorsal could also be phosphorylated after WSSV infection, and we detected Dorsal phosphorylation by using the human p65 Ser276 phosphorylation antibody. The results showed that a strong phosphorylation signal of Dorsal was observed at 3 hpi and 6 hpi (Figure 5E), correlating well with Dorsal enrichment in the nucleus at 3 hpi and 6 hpi, respectively. Taken together, WSSV infection did induce shrimp Dorsal translocated from the cytoplasm to the nucleus, and phosphorylation on Ser342.

**Figure 5.**
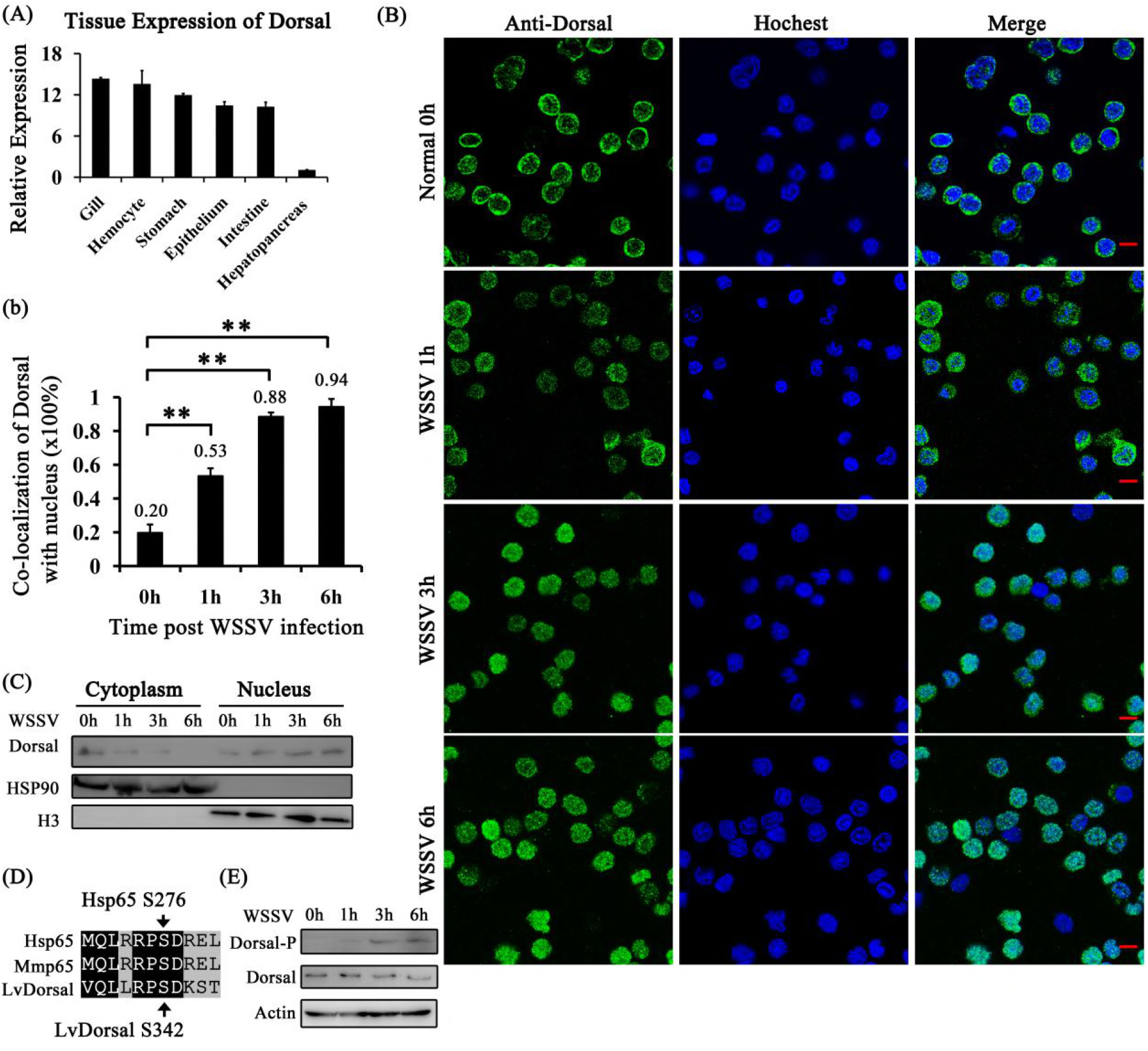
Dorsal nuclear translocation and phosphorylation are induced by WSSV infection. (A) Tissue distribution of Dorsal was analyzed by qRT-PCR. (B) Dorsal nuclear translocation in hemocytes was detected at 1 h, 3 h and 6 h post WSSV infection, and the WSSV untreated hemocytes (0 h) as a control. The hemocytes were collected at 0 h, 1 h, 3 h and 6 h post WSSV infection, deposited onto a glass slide and subjected to immunocytochemical staning by a prepared anti-Dorsal specific antibody, and finally visualized on a fluorescence microscope. (b) Co-localization of Dorsal and Hochest-stained nucleus in hemocytes was calculated by WCIF ImageJ software and analyzed statistically by student’s T test (** *p* < 0.01). (C) The subcellular distribution of Dorsal in hemocytes was detected by western blotting. (D) Dorsal contained a putative phosphorylation site (Ser342) in a conserved region across species. (E) Dorsal was phosphorylated at Ser342 after WSSV infection and analyzed by western blotting with human anti-NF-κB p65 (phospho S276) antibody. All experiments were performed three times, and similar results were obtained.

### Dorsal regulates Toll4 dependent AMP genes expression after WSSV infection

Because Dorsal can be activated upon WSSV infection, we reason that this activation of Dorsal could lead to trigger the expression of some AMPs. To address this, we detected the expression levels of Toll4 dependent AMP genes of ALF1-4 and LYZ1-4 *in vivo* either when Dorsal activity was suppressed by NF-κB inhibitor or when Dorsal expression was silenced by RNAi, respectively. Firstly, we measured whether a NF-κB inhibitor QNZ (EVP4593) can work for Dorsal activity suppression. We observed that the 2 ug/per shrimp was able to suppress Dorsal Ser342 phosphorylation completely *in vivo* and this quality was used in the following analysis (Figure 6A). In order to confirm whether Dorsal regulated AMPs expression, we injected the shrimp with the NF-κB inhibitor QNZ prior to the WSSV challenge, and investigated Dorsal translocation, phosphorylation and AMPs expression. We analyzed the effect of QNZ on Dorsal nuclear translocation in hemocytes under WSSV infection by immunofluorescence staining and western blotting (WB) analysis. The results showed that QNZ significantly suppressed Dorsal translocated from the cytoplasm to the nucleus (Figures 6B and b), which was consistent with the WB analysis of Dorsal cytoplasmic and nuclear localization (Figure 6C). Expectedly, Dorsal Ser342 phosphorylation was efficiently suppressed by QNZ, although shrimps hemocytes were under WSSV challenge (Figure 6D). Further, in the NF-κB inhibitor-injected shrimp the WSSV challenge failed to up-regulate Toll4 dependent AMPs expression both in hemocytes and gills, respectively (Figures 6E and 6F). On the other hand, we carried out RNAi to knockdown Dorsal *in vivo*, WB and qRT-PCR analysis confirmed efficient silencing of its protein levels (Figures 6G) and mRNA levels ((Figures 6H), respectively. We observed that in the RNAi treated shrimps Toll4 dependent AMPs of ALF and LYZ families were marginally down-regulated after WSSV infection compared to GFP dsRNA treated control shrimps both in hemocytes and gills (Figures 6I and 6J). These results persuasively confirm that Dorsal nuclear translocation and phosphorylation are functionally related to the increased expression of Toll4 dependent AMPs under WSSV infection *in vivo*.

**Figure 6.**
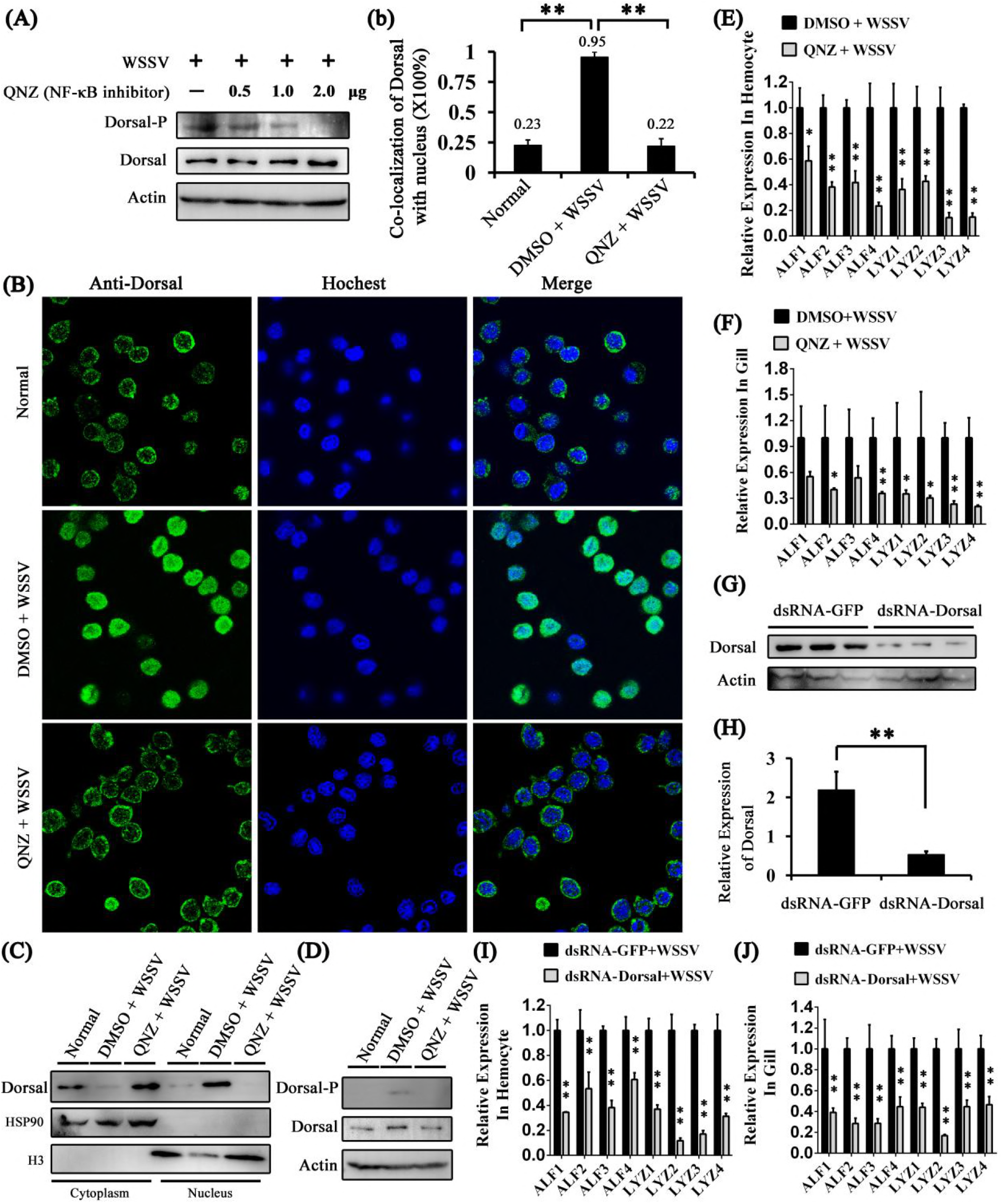
Dorsal regulates the same AMPs modulated by Toll4 upon WSSV infection. (A) Expression of Dorsal-P and Dorsal in QNZ-treated hemocytes at 6 h post WSSV infection were detected by western blotting. (B) Dorsal nuclear translocation in response to WSSV infection was inhibited by QNZ. Each shrimp was injected with 2 μg QNZ (EVP4593), followed by shrimp was intraperitoneal injected with WSSV. The hemocytes were collected at 6 h post WSSV infection, and then subjected to immunofluorescence staining. (b) Co-localization of Dorsal and nucleus in hemocytes corresponding to figure 6B was calculated by WCIF ImageJ software and analyzed statistically by student’s T test (** *p* < 0.01). (C-D) The subcellular distribution (C) and phosphorylation level (D) of Dorsal in QNZ-treated hemocytes was detected at 6 h post WSSV infection by western blotting. The DMSO and WSSV untreated hemocytes were used as controls, respectively. (E-F) AMPs expression levels in hemocyte (E) and gill (F) of QNZ-treated shrimp at 6 h post WSSV infection were detected by qRT-PCR. (G-H) Silencing efficiencies for Dorsal protein levels (G) and mRNA levels (H) were affirmed by western blotting and qRT-PCR. (I-J) AMPs expression levels in hemocyte (I) and gill (J) of Dorsal silenced shrimp at 6 h post WSSV infection were assessed by qRT-PCR. All experiments were performed three times, and similar results were obtained. All the data from b, E, F, H, I and J were analyzed statistically by student’s T test (* *p* < 0.05; ** *p* < 0.01).

### WSSV but not other pathogens-induced Dorsal activation is partially dependent on Toll4 in shrimp

Because both Toll4 and Dorsal induced the same AMPs expression under WSSV infection, we tested whether WSSV mediated Dorsal activation is dependent on Toll4. To assess this, RNAi was performed to evaluate the effects of Toll4 on Dorsal nuclear translocation and phosphorylation under WSSV infection *in vivo*. Efficient silencing of Toll4 mRNA was confirmed by quantitative RT-PCR (Figure 7A). At 6 hours after WSSV infection, but not the negative control (WSSV untreated, Figure 7B top panel), we were able to observe more nuclear imports of Dorsal in hemocytes of GFP dsRNA treated shrimps (96%) than those of Toll4 silenced shrimps (57%) (Figures 7B and b, p < 0.01). We confirmed these results by measuring Dorsal localization and phosphorylation after WSSV infection in hemocytes of GFP dsRNA and Toll4 dsRNA treated shrimps using WB analysis. A decreased percentage of nuclear translocation of Dorsal (46.18%) was observed in Toll4 silenced shrimps under WSSV infection compared to that of dsRNA GFP treated shrimps (86.20%) (Figure 7C and c, p < 0.01). Moreover, knockdown of Toll4 markedly reduced the WSSV induced phosphorylation of Dorsal when compared to GFP dsRNA treated shrimps (Figure 7D and d, p < 0.01). We next appraised the protein levels of shrimp Cactus, an inhibitor of shrimp Dorsal, by WB analysis using an *L. vannamei* Cactus specific antibody. The results demonstrated that a very strong signal and a weak signal were detected in the negative control (WSSV untreated) and Toll4 dsRNA treated shrimps, respectively, but we failed to detect any signal of Cactus in the GFP dsRNA treated shrimps (Figures 7E and e, p < 0.01), which further confirmed that WSSV induced Dorsal activation was partially blocked in Toll4 silenced shrimps. To test whether Toll4 mediated Dorsal activation is pathogen specific, we tested its requirement to other shrimp pathogens including DNA viruses (infectious hypodemal and hematopoietic necrosis virus, IHHNV, and shrimp hemocyte iridescent virus, SHIV), RNA virus (yellow head virus, YHV) and bacteria (*V. parahaemolyticus*). The results showed that all the four types of pathogen could induce Dorsal activation with different degrees of nucleus translocation, but it seems to be not relevant in the context of the silencing of Toll4 (Figure 8). The prominent nuclear translocation of Dorsal in hemocytes after *V. parahaemolyticus* challenge is in good concurrence with previous reports that G^-^ bacterial infection can induce the activation of Dorsal and its translocation (42, 46). These analyses with different pathogens strongly suggest that Toll4 mediated Dorsal activation is WSSV specific, which indicate that Toll4 could play a vital role in recognizing WSSV infection. Taken together, our data suggest that Toll4 is a key factor for sensing WSSV and mediating downstream Dorsal activation, and thereby inducing some specific AMPs production.

**Figure 7.**
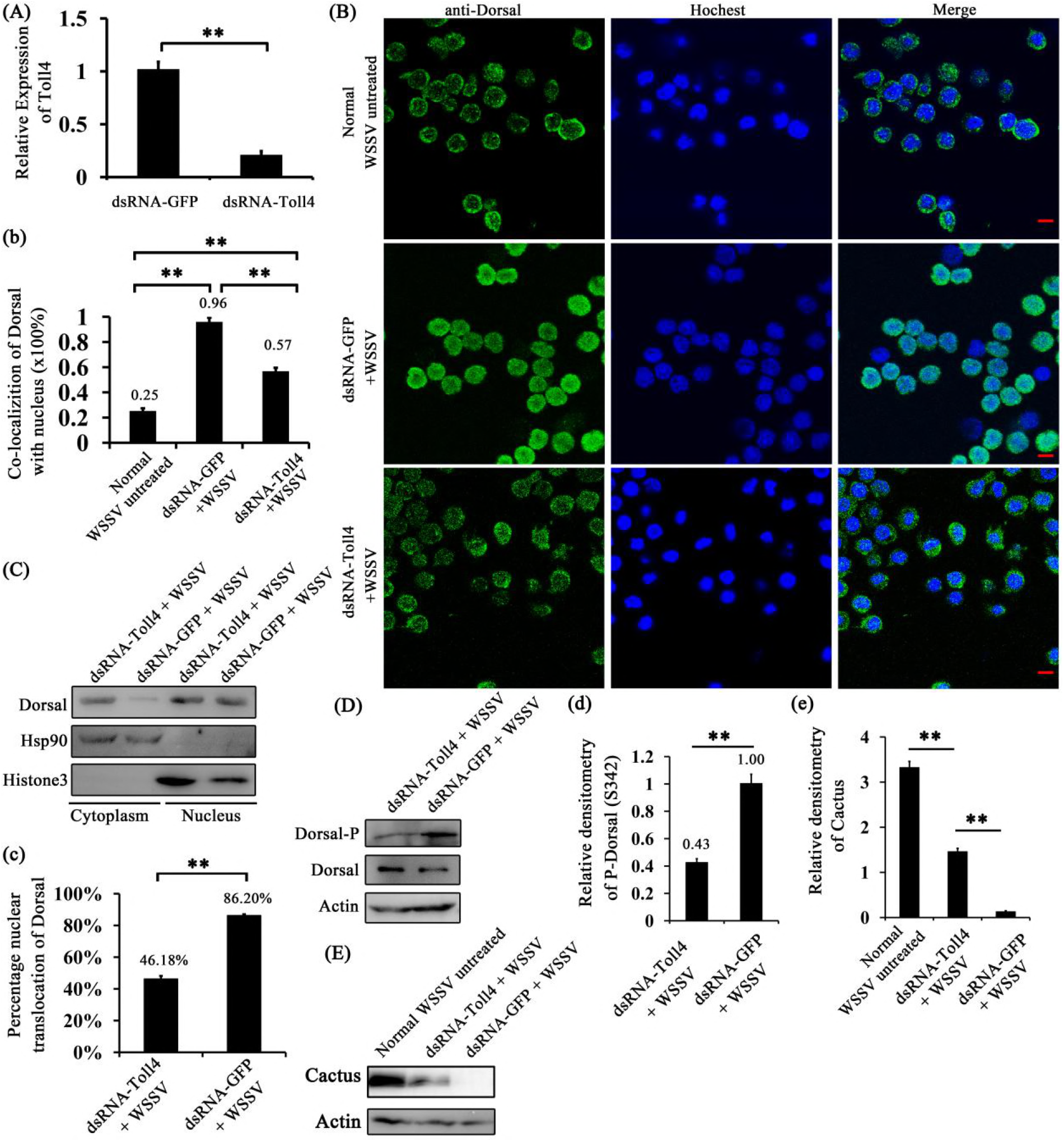
Toll4 regulates Dorsal activation in response to WSSV infection in shrimp. (A) Silencing efficiency for Toll4 in hemocytes was detected by qRT-PCR. (B) Dorsal nuclear translocation in Toll4 silenced hemocytes at 6 h post WSSV infection. (b) Statistical analysis of co-localization of Dorsal and nucleus in hemocytes by WCIF ImageJ software. (C) The subcellular distribution of Dorsal in Toll4 silenced hemocytes was detected at 6 h post WSSV infection by western blotting. The GFP dsRNA treated hemocytes was used as a control. (D) Dorsal phosphorylation level (Ser342) in Toll4 dsRNA and GFP dsRNA treated hemocytes was probed by western blotting. (E) Cactus protein levels were detected by western blotting using prepared anti-Cactus specific antibody in Toll4 dsRNA and GFP dsRNA treated hemocytes at 6 h post WSSV infection. The dsRNA untreated and WSSV non-challenged hemocytes were used as control. (c, d, e) Statistical analysis of the Dorsal nuclear translocation proportion, Dorsal phosphorylation levels and Cactus protein levels in Toll4 dsRNA and GFP dsRNA treated hemocytes at 6 h post WSSV infection, respectively. All experiments were performed three times, and similar results were observed. All the data from b, c, d and e were analyzed statistically by student’s T test (** *p* < 0.01).

**Figure 8.**
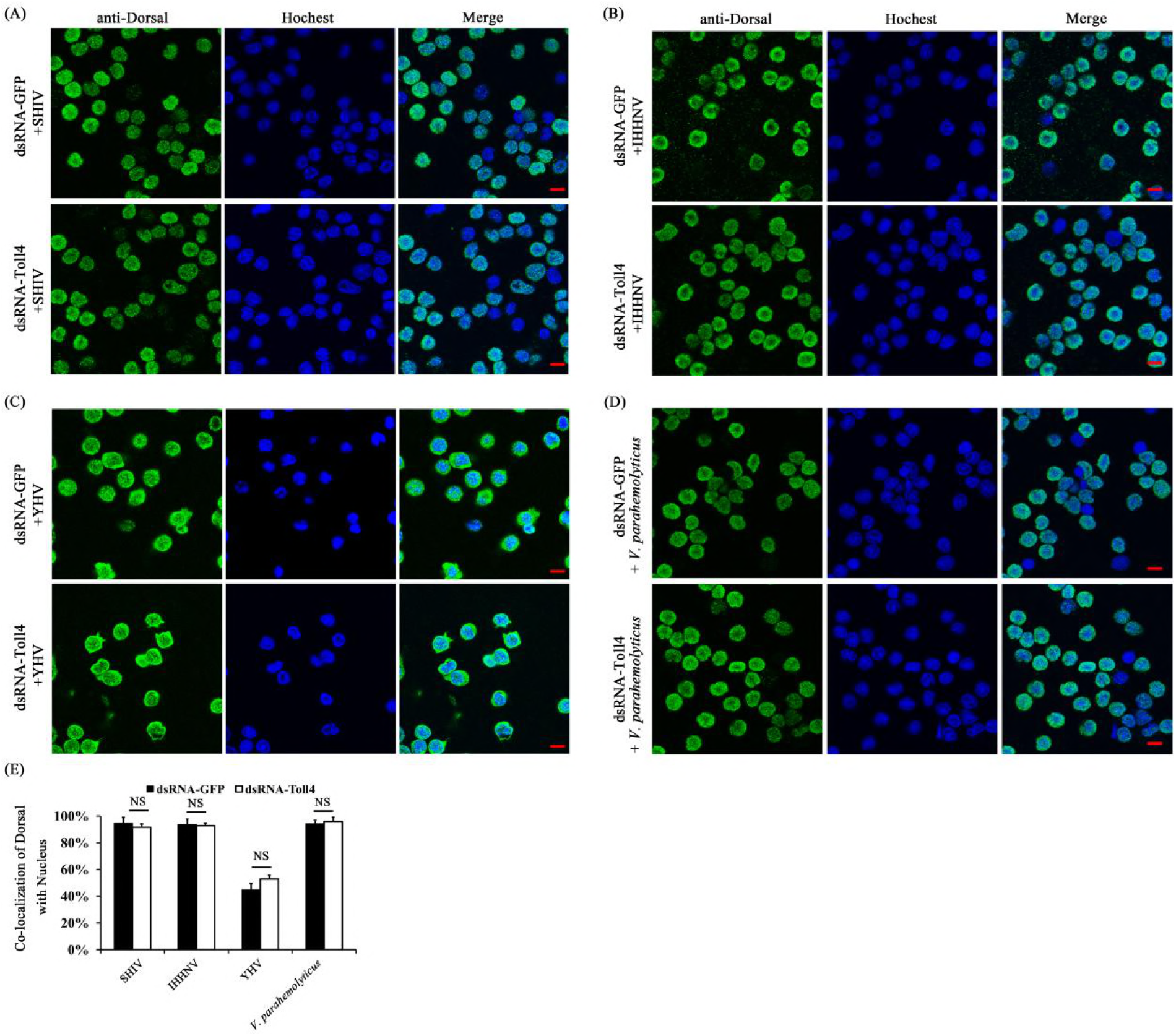
Knockdown of Toll4 doesn’t affect the translocation of Dorsal in response to other shrimp pathogens. (A-D) The immunocytochemical staning of Dorsal in Toll4 silenced hemocytes with challenges of SHIV (A), IHHNV (B), YHV (C) and *V. parahaemolyticus* (D) at 6 hpi as described earlier. The GFP dsRNA treated hemocytes following infection were used as a control. (E) Co-localization of Dorsal and Hochest-stained nucleus in hemocytes were calculated by WCIF ImageJ software and analyzed statistically by student’s T test (NS, not significant).

### AMPs regulated by Toll4-Dorsal pathway oppose WSSV infection

The induction of AMPs as a response to pathogenic infection is a crucial defense mechanism of innate immunity in invertebrates including insects and shrimps. Our results above have revealed that after WSSV infection Toll4 and Dorsal induced the same AMPs expression, raising the hypothesis of these AMPs playing antiviral role against WSSV. We silenced the eight AMPs including ALF1-4 and LYZ1-4 regulated by both Toll4 and Dorsal through dsRNA treatment. Silence efficiencies were confirmed by quantitative RT-PCR (Figure 9A). To gain more information about the effects of AMPs on viral replication, a timeframe of 24 h, 48 h, 72 h, 96 h and 120 h after WSSV infection was selected to investigate the viral loads in gills of each AMP silenced shrimp. Compared with GFP dsRNA inoculated shrimps, shrimps in which ALF1, ALF2, ALF3, ALF4, LYZ1, LYZ2, LYZ3 or LYZ4 were silenced had higher viral burden (Figures 9B, 9C, 9D, 9E and 9F) at the whole timeframe. To further dissect the function of these AMPs during WSSV infection, a parallel experiment was performed to explore the survival phenotype of each AMP silenced shrimps followed by WSSV infection. Experimental shrimps were challenged with WSSV at 48 h post dsRNA injection, and the survival rate was recorded over a period of 168 h after the challenge. We observed that in the each AMP-knockdown group except LYZ3-knockdown group, shrimps were more susceptible to WSSV infection (Figures 9G and 9H). Notably, the survival rate of LYZ3-knockdown group showed no significant difference, but had the trend of lower (p = 0.3853) compared to that of control group (Figure 9H). In summary, our data convincingly demonstrate that Toll4-Dorsal pathway regulated AMPs are involved in WSSV restriction in shrimp.

**Figure 9.**
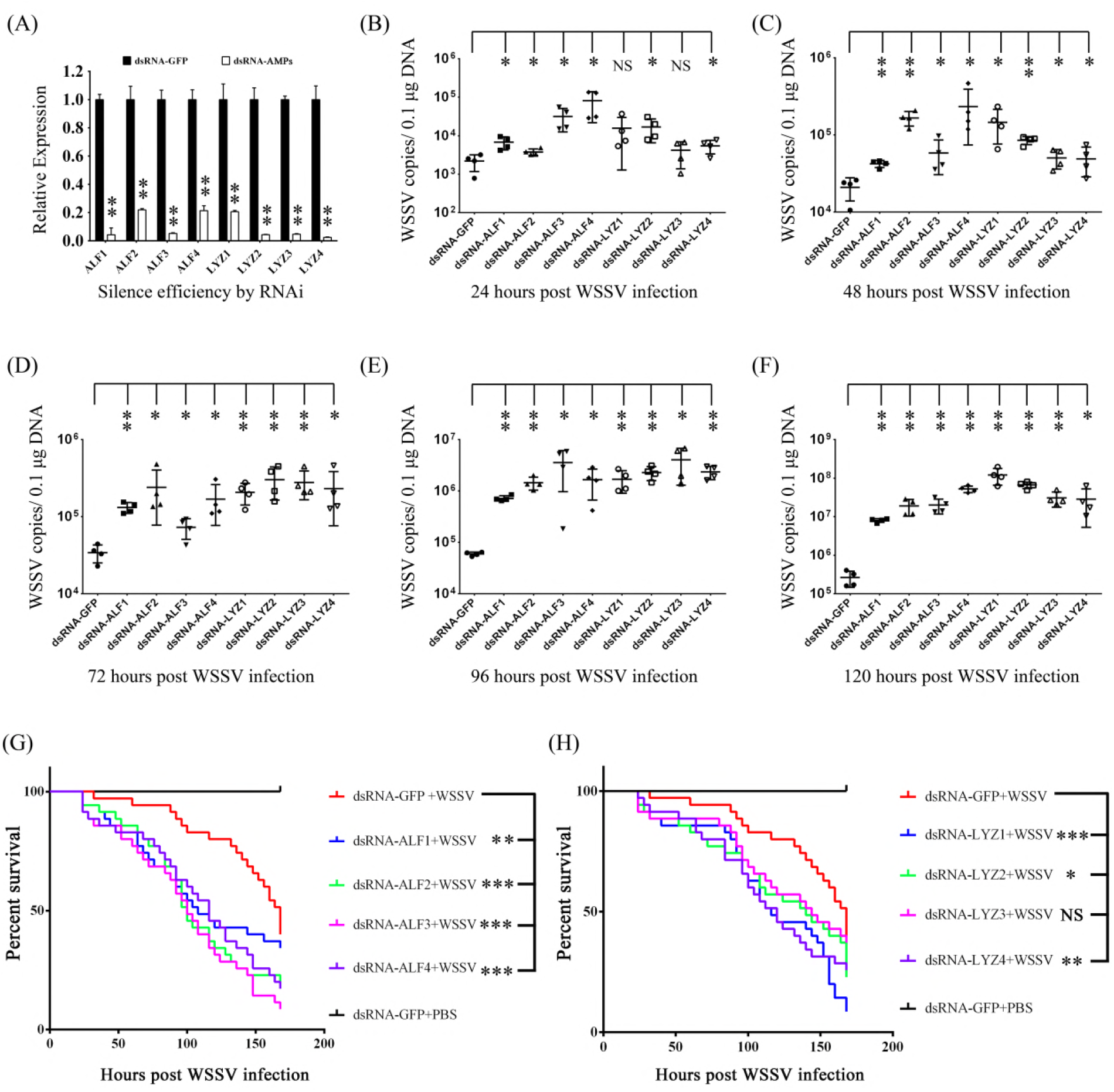
The function of Toll4-Dorsal cascade regulated AMPs in WSSV infection. (A) Effective knockdown for ALF1-4 and LYZ1-4 in hemocytes by dsRNA was confirmed by qRT-PCR. (B-F) Silencing of AMPs enhanced WSSV infection in shrimp. WSSV was inoculated at 48 h post each AMP silencing. The viral loads in gills were assessed at 24 h (B), 48 h (C), 72 h (D), 96 h (E) and 120 h (F) post-infection via absolute qPCR. (G-H) Survival of WSSV challenged AMP-silenced shrimp and GFP dsRNA treated shrimp. All experiments were performed three times, and similar results were obtained. All the data from B, C, D, E and F were analyzed statistically by student’s T test (* *p* < 0.05; ** *p* < 0.01). Survival rates from G and H were analyzed statistically by the Kaplan–Meier plot (log-rank χ^2^ test) (** *p* < 0.05, ** *p* < 0.01, NS, not significant).

### AMPs exhibit antiviral activity by interacting with WSSV structural proteins

Some AMPs have been reported to play a vital role in combating viral pathogens via directly acting on the viral virion (47, 48). To decipher the molecular basis underlying AMPs against WSSV, we performed an *in vitro* pull-down assay between AMPs and WSSV structural proteins in order to elucidate how these AMPs act on WSSV. In this study, we pay our attention to ALF1 and LYZ1 as a representative one of ALF and LYZ families, respectively. Six WSSV structural proteins including VP19, VP24, VP26, VP28, wsv134 and wsv321 were used in the *in vitro* pull-down assay to explore the potential interaction between the above-mentioned structural proteins and ALF1 or LYZ1. The six WSSV structural proteins with GST tag and the two AMPs ALF1 and LYZ1 with His tag were expressed and purified (Figures 10A and 10B). In the His tagged ALF1 pull-down assay with six WSSV structural proteins (GST tag), we observed that ALF1 precipitated VP19, VP26, VP28, wsv134 and wsv321 by SDS–PAGE with coomassie blue staining (Figure 10C, lanes 1, 3, 4, 5 and 6, respectively). However, His tagged ALF1 did not interact with GST tagged VP24, which indicates that the interaction of between ALF1 and other four structural proteins is specific, but not related to the His and GST tags. We further confirmed this result by western blotting with GST tag antibody, which is in good agreement with that of coomassie blue staining (Figure 10D). In the His tagged LYZ1 pull-down assay, we found that VP26, VP28, wsv134 and wsv321 were enriched (Figure 10E, lanes 3, 4, 5 and 6, respectively), and an identical result was observed by western blotting (Figure 10F). To further identify the above results, six WSSV structural proteins with GST tag was used in a GST pull-down assay with purified His tagged ALF1 or LYZ1 followed by SDS-PAGE with coomassie blue staining and western blotting with His antibody, respectively. As shown in Figures 10G and 10H, VP19, VP26, VP28, wsv134 and wsv321 interacted with ALF1 (arrows) in GST pull-down assay, which further confirmed the results of pull-down assay with His tagged ALF. In a similar manner, Figures 10I and 10J showed that VP26, VP28, wsv134 and wsv321 strongly interacted with LYZ1. Thus, the results strongly suggest that ALF1 and LYZ1 were able to interact with WSSV structural proteins, to be specific, ALF1 interacted with VP19, VP26, VP28, wsv134 and wsv321, and LYZ1 interacted with VP26, VP28, wsv134 and wsv321 (Figure 10K).

**Figure 10.**
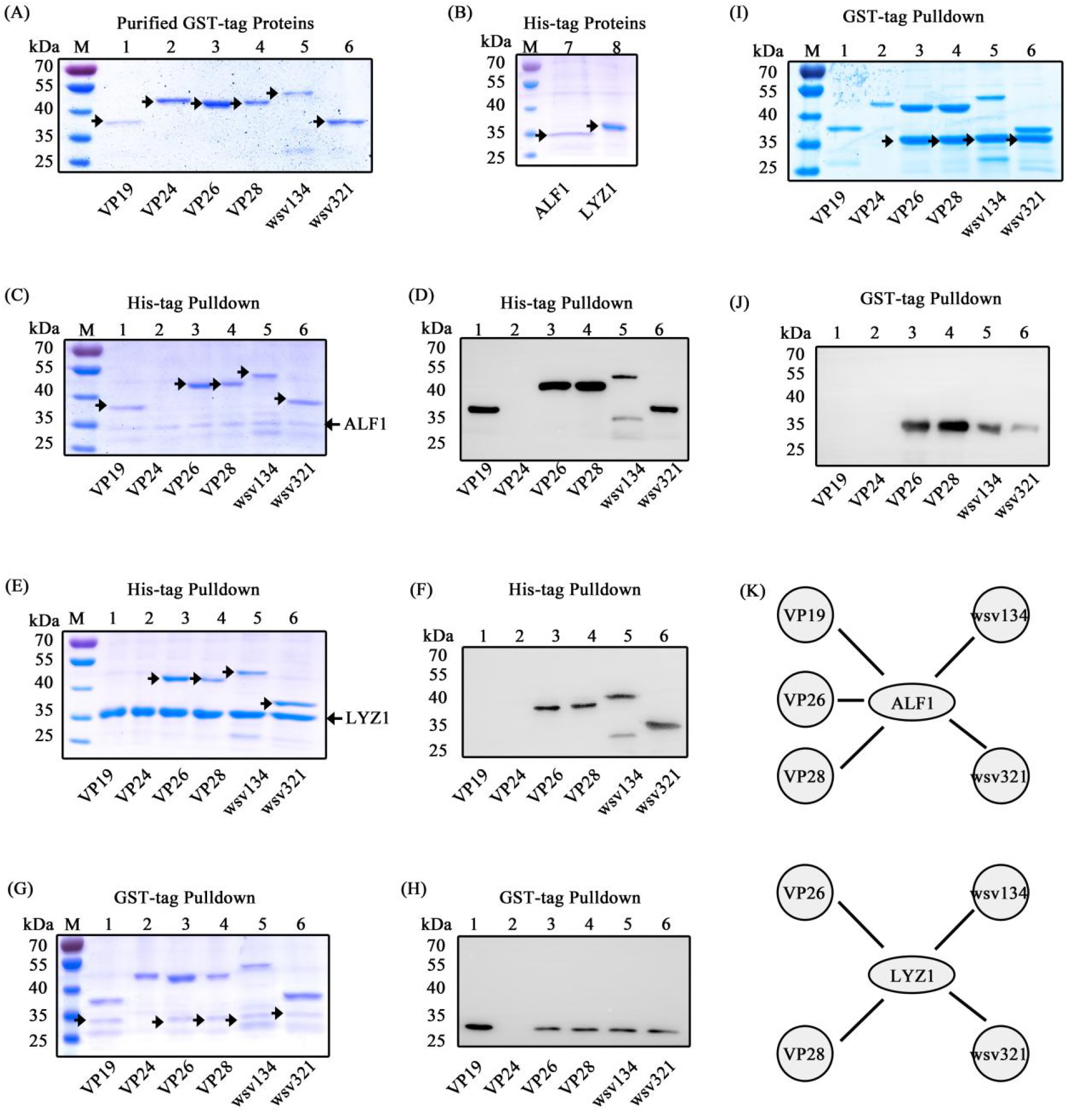
AMPs interact with WSSV structural proteins. (A) Purified GST tagged WSSV structural proteins of VP19, VP24, VP26, VP28, wsv134 and wsv321. (B) Purified His tagged ALF1 and LYZ1. (C) His tagged ALF1 interacted with GST-VP19, VP26, VP28, wsv134 and wsv321 was obtained in His pull-down assay and visualized by coomassie blue staining. (D) His-ALF1 interacted with GST-VP19, VP26, VP28, wsv134 and wsv321 in His pull-down assay was confirmed by western blotting with anti-GST antibody. (E) His tagged LYZ1 interacted with GST tagged VP26, VP28, wsv134 and wsv321 was obtained in His pull-down assay and visualized by coomassie blue staining. (F) His pull-down assay with His-LYZ was confirmed by western blotting with anti-GST antibody. (G) GST tagged VP19, VP24, VP26, VP28, wsv134 and wsv321 were used to pull down His-ALF1, and visualized by coomassie blue staining. (H) GST pull-down assay of GST-VP19, VP26, VP28, wsv134 and wsv321 interacting with His-ALF1 was confirmed by western blotting with anti-His antibody. (I) GST tagged VP19, VP24, VP26, VP28, wsv134 and wsv321 were used to pull down His-LYZ1, and visualized by coomassie blue staining. (J) GST pull-down assay of VP26, VP28, wsv134 and wsv321 interacting with His-LYZ1 was confirmed by western blotting with anti-His antibody. (K) Schematic illustration of the ALF1 or LYZ1 interacting with WSSV structural proteins. The experiments were repeated three times.

## Discussion

Accumulating evidence indicates that shrimp Tolls participate in host defense against WSSV infection; however, the underlying mechanism of the Toll receptor mediated antiviral functions has been poorly understood. Herein, we have identified an antiviral role for a new Toll from *L. vannamei*, the Toll4, in response to WSSV infection *in vivo*. Toll4 silenced shrimps demonstrate significantly elevated viral replication and mortality after WSSV challenge. Shrimps with knockdown of genes in some core components of the canonical Toll pathway such as MyD88, Tube, Pelle and Dorsal have remarkably increased WSSV titers. Furthermore, Toll4 appears to be specific to sense WSSV infection to trigger Dorsal, which lead to induce a specific set of AMPs with the ability of interacting with viral structural proteins that confer resistance to viral infection. Our results have now demonstrated that the Toll4-Dorsal-AMPs cascade is involved in the control of WSSV infection in shrimp.

The Toll pathway is essential to establish an innate immune response to defend against a wide range of pathogens including virus. The importance of this pathway in the innate control of viral infections in insects is best demonstrated by that mutants in some core components such as Dif and Toll of *Drosophila* with increased susceptibility to infection (10). The *Drosophila* Toll pathway has also been shown to play an universal antiviral role against multiply viruses by oral infection such as *Drosophila* C virus, Cricket paralysis virus, Flock house virus, and Nora virus (49). In addition, several reports show that the Toll pathway has an antiviral role in innate immunity of mosquitoes (50-52). Our results indicate that the increased lethality rates observed in the Toll4 silenced shrimps are associated with higher WSSV loads. Thus, the Toll4 is involved in resistance to WSSV and it is a major antiviral factor in shrimp. Moreover, several Tolls have been proved to confer antiviral immunity in other shrimps. For example, a Toll4 from *P. clarkii* (29) and a Toll from *M. rosenbergii* (31) are important for the innate immune responses against WSSV, although the exact antiviral mechanism is not elucidated. We also provide evidence indicating the key antiviral role of the canonical Toll pathway by that silencing of the core components such as MyD88, Tube, Pelle and Dorsal results in increased WSSV titers. These data may suggest that the function of Toll pathway in the control of viral infections could be conserved through evolution. This is consistent with previous studies showing Toll pathway antiviral effect in other Arthropods including crayfish (28), *Drosophila* (10, 49), mosquitoes (53) and honeybees (50).

In general, the canonical Toll pathway of *Drosophila* mediated immune response relies on the activation of the NF-κB transcription factors Dorsal or Dif, however whether it is true for shrimp is still largely unknown. Firstly, we demonstrate that Toll4 and Dorsal are involved in regulating the same AMPs after WSSV infection. In addition, our data show that detection of WSSV infection by Toll4 triggers transcriptional activity of Dorsal, but knockdown of Toll4 was not sufficient to restrain the activation of Dorsal in response to WSSV infection. This may be owing to inability to absolutely suppress Toll4 expression by RNAi method. But we cannot exclude the possibility that the activation of Dorsal in response to WSSV infection may integrate signals from other upstream receptors. In other words, there could be more than one upstream receptor, in addition to Toll4, involved in the response to WSSV infection responsible for the activation of Dorsal. Of note, Dorsal has the ability to bind with the promoters of some WSSV genes such as the Immediate Early gene 1 (IE1) and regulate their transcriptional expression in insect cells background or *in virto* (54). Thus, it is thought that Dorsal is required for WSSV gene expression and genome replication (55), more experiment evidence *in vivo* needs to support this. By RNAi method, we observe that shrimps with knockdown of Dorsal have elevated viral loads than those of the GFP control group, suggesting that Dorsal is important for host to limit viral replication. Besides, the report of WSSV encoding two MicroRNAs with the ability to suppress shrimp Dorsal also supports that Dorsal is a key restrict factor against viral infection (56). However, knockdown of Dorsal results in lower viral loads than those of MyD88, Tube and Pelle silenced groups, which may be explained by that Dorsal locates the lower levels at MyD88/Tube/Pelle cascade of the canonical Toll pathway.

Some reports have showed that Tolls from *Drosophila* and shrimps inducing antiviral innate immunity are independent of activation of the transcription factor NF-κB (Dorsal or Dif), as shown by the fact that *Drosophila* Toll7 activates antiviral autophagy not involvement of Dorsal or Dif (11), as well as by the fact that shrimp Toll3 initiates a IRF-Vago dependent antiviral route (33). Therefore, whether other Tolls are responsible for activation of Dorsal in response to WSSV infection, and how they confer resistance to WSSV infection deserve to be further studied. In addition, in the present study, we identified a total of nine Tolls, and silencing of each of Toll expect for the Toll2 contribute to increased WSSV loads compared to the control shrimps. This observation is reminiscent of that the lack of immunoglobulin-based adaptive immune system and classical IFN mediated antiviral defense maybe require some invertebrates including shrimps and *Drosophila* to be more heavily dependent on the Tolls or other receptors for antiviral immunity. Identification of the target genes of the other Tolls after viral infection will be important to understand how they contribute to resistance to viruses.

There are significant differences in the Toll and TLR receptors initiated activation by ligand recognition in invertebrate and vertebrate. In general, TLR receptors in mammals are able to detect microbial infection through directly binding to PAMP (2), but *Drosophila* Toll1 instead interacts with the endogenous cytokine-like factor Spätzle, the product of a proteolytic cascade induced upon upstream recognition of fungal and bacterial PAMPs (51). Notablely, Toll7 in *Drosophila* can bind to vesicular stomatitis viruses at the plasma membrane and therefore has been considered as a specific and bona fide PRR for sensing this virus (11). Besides, several Tolls from *L. vananmei* are able to detect some PAMPs directly, as shown by the fact that Toll1 and Toll3 can interact with CpG ODN 2395 *in vitro* (57). Surprisingly, another study demonstrates that three Tolls from *M. japonicas*, two of them homologous to the above Toll1 and Toll3 from *L. vananmei* (57), can directly bind to both PGN and LPS (46). These data suggest that one type of shrimp Toll could recognize more than one PAMP. In the present study, we observed that Dorsal activation and translocation to the nucleus is dependent on Toll4 in response to WSSV infection, but not other tested pathogens, which indicate that Toll4 could also be a specific PRR to detect WSSV in a manner similar to *Drosophila* Toll7. Unravelling how Toll4 senses WSSV in the future will be important to understand antiviral immunity in shrimp.

The production of antimicrobial peptides (AMPs) is commonly considered to be an evolutionarily conserved mechanism of the innate immune response and has been extensively studied in vertebrates and other non-vertebrate organisms including shrimps. Some shrimp Tolls are able to resist bacterial infection via regulating a wide range of AMPs expression (28, 29, 58), which inspires us to suppose that shrimp Tolls can also regulate some specific AMPs synthesis to oppose WSSV. In fact, our examinations of AMPs expression in WSSV-infected shrimp show an increase in expression of AMPs comparable to that found during a *V. parahaemolyticus* infection of shrimp. Moreover, the decreased expression of the same AMPs in Toll4 and Dorsal-silenced shrimp does show that Toll4-Dorsal pathway indeed devotes to induce AMPs transcription upon WSSV infection. By RNAi, we detect survival rates and viral titers in single AMPs silenced shrimps and find that each single AMP except for LYZ3 provides effective resistance to viral infection. Although evidence exists that some AMPs can respond to WSSV infection, it was not known how these shrimp AMPs affected viruses. Previous reports show that shrimp ALF can protect against WSSV infection via interfering with viral replication *in vitro* and *in vivo* in crayfish *Pacifastacus leniusculus* (59) and CqALF can disrupt WSSV envelope integrity that leads to the decrease of WSSV infectivity in the red claw crayfish *Cherax quadricarinatus* (60). Furthermore, an ALF isoform 3 from *P. monodon* has performed its anti-WSSV action by binding to several viral structural proteins such as wsv131 (WSSV186), wsv134 (WSSV189) and wsv339 (WSSV395) (61). On the other hand, lysozyme is a key effector of the innate immune system and kills bacteria by catalytic hydrolysis of cell wall peptidoglycan, but it also exhibits catalysis-independent antimicrobial properties. For example, human lysozyme has been shown to inhibit HIV-1 infection *in vitro* by preventing the adsorption and penetration of the virus (62, 63). HL9, a nonapeptide fragment of human lysozyme, blocks HIV-1 viral entrance and replication by binding to the ectodomain of gp41, the envelope glycoprotein of HIV-1 crucial to membrane fusion (63, 64). These data strongly suggest that ALF and LYZ family have effective antiviral activity, and it seems reasonable to hypothesize that shrimp ALF and LYZ family are able to interact with WSSV structural proteins. In this study, one anti-LPS-factor ALF1 and one Lysozyme LYZ1 are chosen to explore their antiviral actions. In agreement with this hypothesis, our results reveal that ALF1 interacts with VP19, VP26, VP28, wsv134 and wsv321, while LYZ1 interacts with VP26, VP28, wsv134 and wsv321. Given their conserved sequences, the other AMPs, in addition to ALF1 and LYZ1, could be able to interact with some specific WSSV structural proteins. Interestingly, currently, no mechanistic analysis on LYZ family genes responsible for antiviral role against WSSV has been performed except for a role in modulating the humoral response to this virus infection (65)). Thus, LYZ1 in this study is the first LYZ family gene identified with the capacity to interfere with replication of this important pathogen, which suggest that LYZ could be a new type of effectors for restricting WSSV infection. Of note, one type of AMP can interact with more than one WSSV structural protein, and vice versa. Likewise, WSSV structural proteins VP19, VP26 and VP28 are shown to interact with each other to form a multiprotein complex (66). So, it seems that AMPs interact with WSSV structural proteins as a manner of multiply layers or reticulation, which could be more effective to control virus. In addition to their important functions in maintaining the integrity of virion, WSSV structural proteins also play a key role in initiating viral infection (67), as showed by fact that some structural proteins such as VP28 and VP26 are shown to being key factors essential for virus attachment and entry into host cells (67-70). Therefore, based on our data together with previous reports, it is highly conceivable that the Toll4-Dorsal pathway regulated AMPs interacts with WSSV structural proteins to both disrupt WSSV integrity and interfere with viral invasion.

In summary, we clone and identify a total of nine Tolls from *L. vannamei*, RNAi screens Toll4 as a key antiviral factor against WSSV infection. Considering the data obtained in the present study, we propose the following antiviral immune signaling pathway in *L. vannamei* (Figure 11): i. viral recognition and signal transduction: Toll4 recognizes WSSV infection to converge on Dorsal translocated from the cytoplasm to the nucleus and phosphorylation on Ser342; ii. AMP induction: the activation of Dorsal in the nucleus triggers a specific set of AMPs expression such as ALFs and LYZs; and iii. viral inactivation: Toll4-Dorsal drived AMPs can bind with the components of the viral surface, subsequently resulting in the WSSV inactivation. Uncovering the Toll4-Dorsal-AMPs cascade mediated antiviral program may provide novel strategies for limiting WSSV infection in shrimp aquaculture, and dissecting the pattern of Toll4 sensing WSSV in the further will provide additional insights into how the canonical Toll pathway responds to viral infection.

**Figure 11.**
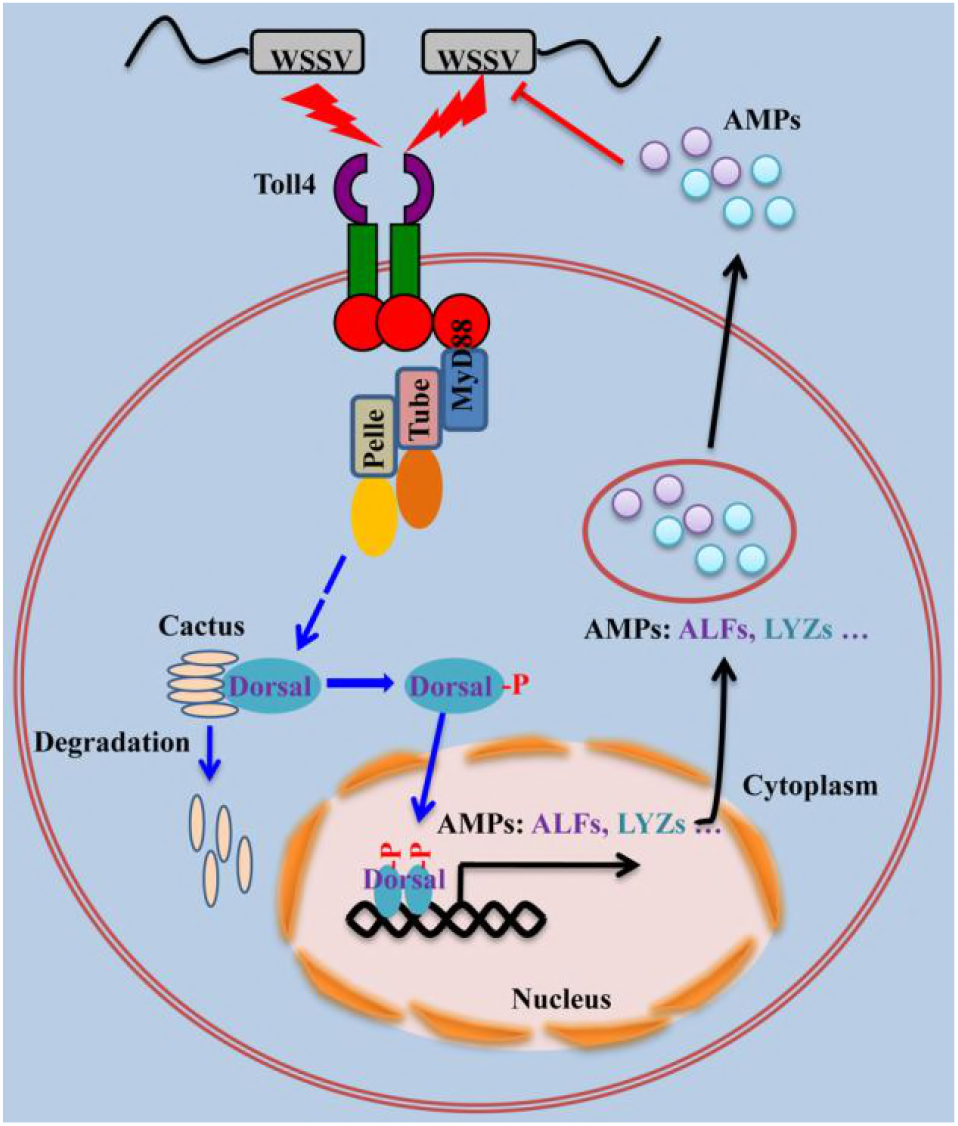
Model for Toll4 mediated antiviral mechanism against WSSV. In shrimp, Toll4 sensed WSSV infection and proceeded to the degradation of shrimp IκB factor Cactus. After Cactus degradation, the transcription factor Dorsal was phosphorylated and translocated into the nucleus, where it led to activation of the transcription of several sets of AMPs (ALF1-4 and LYZ1-4). These effector molecules (AMPs) were secreted to extracellular space and executed anti-WSSV activity through interacting with its structural proteins.

## Materials and methods

### Animals and pathogens

Shrimps (*L. vannamei*, average weight 8 g each) were purchased from the local shrimp farm in Zhuhai, Guangdong Province, China, and fed with a commercial diet in a recirculating water tank system filled with air-pumped sea water (2.5% salinity) at 28 °C. Before all experiment treatments, the shrimps (5% of total) were detected and confirmed to be free of common pathogens including white spot syndrome virus (WSSV), yellow head virus (YHV), taura syndrome virus (TSV), shrimp hemocyte iridescent virus (SHIV, also known as CQIV), infectious hypodemal and hematopoietic necrosis virus (IHHNV) and *Vibrio parahaemolyticus* by PCR or RT-PCR methods according to standard operation procedures by Panichareon et al (71) and Qiu et al (72). Because many genes from the shrimp canonical Toll-Dorsal pathway can be activated by Gram-negative (G^-^) bacteria (42, 46), *V. parahaemolyticus* thus was used here as a positive activator of the shrimp Toll-Dorsal pathway. The Gram-negative bacteria *V. parahaemolyticus* were cultured in Luria broth (LB) medium overnight at 37 °C, and the bacteria were harvested by centrifugation (5000 g, 10 min) and washed twice in phosphate buffer saline (PBS) to remove growth medium and finally resuspended in PBS to give 10^8^ cells per ml. A final injection density of *V. parahaemolyticus* was adjusted to yield approximately 1 × 10^5^ CFU/50 μl as a previous study (73). WSSV was extracted from the WSSV-infected shrimp muscle tissue and stored at −80 °C. Before injection, muscle tissue from WSSV infected shrimp was homogenized and prepared as WSSV inoculum with approximately 1 × 10^5^ copies in 50 μl PBS following a published method (74). In the pathogenic challenge experiments, each shrimp was received an intraperitoneal injection of 50 μl WSSV or *V. parahaemolyticus* solution at the second abdominal segment by a 1-ml syringe.

### Cloning of shrimp Tolls

In order to obtain the cDNA sequence of all candidate Toll genes from shrimp, the amino acid (aa) sequences of the Tolls and TLRs from *Drosophila* and human (DmToll1-9 and HsTLR1-10) were collected and used as query sequences for in silico searches of *L. vannamei* transcriptome data (34) using local TBLASTN alignment tool with E-value cutoff of 1e^−5^. Nine assembled EST sequences were identified as having high homology to the Toll family genes. Gene-specific primers (Supplement Table 1) were designed for 5′ and 3′ rapid amplification of cDNA ends (RACE) PCR to obtain the 5′ and 3′ end of *L. vannamei* Toll genes. In brief, total RNA was extracted from pooled tissues of *L. vannamei* gill, hemocyte and intestine followed by the protocol described in the RNeasy Mini Kit (Qiagen). cDNA synthesis, 5′/3′-rapid amplification of cDNA ends (5′/3′-RACE) PCR, and nested PCR were performed using a SMARTer RACE cDNA amplification kit (Clontech, Japan) in accordance with the manufacturer’s instructions. The final PCR products were cloned into pMD-19T Cloning Vector (TaKaRa, Japan) and 12 positive clones were selected and sequenced. Then, we performed TBLASTN again by using the aa sequences of nine Tolls as query sequences to search against several RNA-Seq databases from NCBI and others (35-38), but no new Toll was identified, suggesting there could be just a total of nine Tolls in shrimp.

### Sequence and phylogenetic analysis

Protein domains of Tolls and TLRs were identified by using Simple Modular Architecture Research Tool (SMART) (http://smart.embl.de/). Shrimp TIR domains of nine Tolls were aligned by using Clustal X v2.0 program (75) and GeneDoc software where the identities among each other were labeled. The neighbor-joining (NJ) phylogenic tree was constructed based on the deduced amino acid sequences by utilizing MEGA 5.0 software (76).

### Antibodies

The polyclonal antibodies for *L. vannamei* Dorsal and Cactus were produced in guinea pigs and rabbits, respectively, by GL Biochem antibody manufacturing company (China) from our previous study (44). Polyclonal rabbit anti-NF-κB p65 (phospho S276) antibody (ab194726, Abcam) was used to detect the phosphorylated shrimp Dorsal. Rabbit anti-Histone H3 (4499s), Rabbit anti-Hsp90 (ab13495), and the secondary antibodies Goat Anti-Guinea pig IgG H&L (Alexa Fluor^®^ 488) (ab150185), Goat anti-Guinea pig IgG H&L (HRP) (ab6908), anti-Mouse IgG H&L (HRP) (ab6789) and anti-Rabbit IgG H&L (HRP) (ab6721), were purchased from Abcam (USA). Mouse anti-Actin antibody was obtained from Merck (MAB1501). Mouse anti-6His antibody (H1029) and Mouse anti-GST antibody (SAB4200237) were gained from Sigma (USA).

### Detection of viral loads by absolute quantitative PCR

In order to monitor the WSSV copies, absolute quantitative PCR (qPCR) was conducted by utilizing with the forward and reverse primers of wsv069 (WSSV32678-F/WSSV32753-R), a WSSV single copy gene, and a TaqMan fluorogenic probe (WSSV32706) followed by a published method (55). The primers used here were shown in Supplement Table 1. In brief, a 675-bp DNA amplicon of wsv069 with a region of 32678 to 32753 from WSSV genome (AF332093.2) was obtained and subcloned into the pMD19-T plasmid. The plasmid pMD19-T containing the 675-bp DNA fragment was used as the internal standard, and serially diluted to 10-folds to generate a standard curve of absolute qPCR. Genomic DNA from shrimp muscle, hepatopancreases and/ or gill was extracted with Marine Animal Tissue Genomic DNA Extraction Kit (TianGen Biochemical Technology). The extracted shrimp DNA and the internal standard plasmid were subjected to absolute qPCR. The PCR reaction mixture and cycling conditions were the same as previous research (55). Each sample from one shrimp was made in three replicates by absolute qPCR. The WSSV genome copies were calculated and normalized to 0.1 μg of shrimp tissue DNA.

### Semi-quantitative and quantitative reverse transcription PCR

Semi-quantitative reverse transcription PCR (Semi-qRT-PCR) was used to analysis the tissue distribution of nine Tolls in uninfected shrimp. Briefly, healthy shrimp tissues including hepatopancreases, gill, intestine, hemocyte, stomach, epithelium, heart and muscle were sampled. Three samples from each tissue were collected from 15 shrimps (5 shrimps pooled together). Total RNA was extracted from each tissue with RNeasy Mini Kit (Qiagen), and reverse transcribed to cDNA with PrimeScript™ II 1st Strand cDNA Synthesis Kit (Takara) following the manufacturer’s instructions. The cDNA fragments of nine Tolls were amplified using the gene specific primers (Supplement Table 1) under the following conditions: 1 cycle of 94 °C for 2 min, 28 cycles of 94 °C for 30 s, 60 °C for 30 s, 72 °C for 30 s, followed by elongation at 72 °C for 5 min. As an internal loading control, the shrimp EF1α (GU136229) was amplified as the same PCR conditions.

Quantitative reverse transcription PCR (qRT-PCR) was conducted to detect the mRNA levels of genes (Tolls, Toll pathway components or AMPs) under the pathogenic challenge experiments or RNAi *in vivo*. The method of tissues collection, total RNA extraction and cDNA synthesis was as described above. qRT-PCR analysis was performed in the LightCycler 480 System (Roche, Germany) with a volume of 10 μl comprised of 1 μl of 1:10 cDNA diluted with ddH_2_O, 5 μl of 2 × SYBR Green Master Mix (Takara, Japan), and 250 nM of each primer (Supplement Table 1). The cycling programs were the following parameters: 95 °C for 2 min to activate the polymerase, followed by 40 cycles of 95 °C for 15 s, 62 °C for 1 min, and 70 °C for 1 s. Cycling ended at 95 °C with 5 °C/s calefactive velocity to create the melting curve. Expression level of each gene was calculated relative to internal control gene EF-1α by using the Livak (2^-ΔΔCT^) method.

### DsRNA production and RNAi performance

The dsRNAs including Dorsal (accession No. ACZ98167), Tube (KC346865), Pelle (KC346864), MyD88 (AFP49302), nine Tolls (Supplement data S1), eight AMPs (Supplement data S2) and the control GFP, were synthesized by T7 RiboMAX™ Express RNAi System kit (Promega, USA) followed by the user’s manual. More detailed information about the primers for dsRNA synthesis was listed in Supplement Table 1. The quality of dsRNA was checked after annealing via gel electrophoresis. The RNA interference (RNAi) assay was performed as we described else (73). Briefly, each shrimp was received an intraperitoneal injection at the second abdominal segment of dsRNAs (20 μg) for Dorsal, Tube, Pelle, Toll1, Toll2, Toll3, Toll4, Toll5, Toll6, Toll7, Toll8, Toll9, ALF1, ALF2, ALF3, ALF4, LYZ1, LYZ2, LYZ3, LYZ4 or GFP (as a control). The gills and/ or hemocytes were collected from the shrimp 48 h after the dsRNA injection, and total RNA was extracted and assessed by qRT-PCR using the corresponding primers (Supplement Table 1) to evaluate the efficacy of RNAi.

To screen potential Toll with antiviral effects against WSSV, shrimps were divided into ten groups: one control group received GFP dsRNA injection and the other nine RNAi groups received each Toll dsRNA injection, respectively. At 48 h after the RNAi performance, each shrimp was challenged with 10^5^ copies of WSSV particles by intraperitoneal injection, and 48 hours later again, muscle, hepatopancreas and/ or gill tissue from 12 shrimps was sampled to examine the virus copies by absolute qPCR. To further investigate whether the lethality rates of Toll4 silenced shrimps was associated with viral levels, hepatopancreas, gill and muscle was sampled at 48 hpi to examine the virus copies by absolute qPCR. To evaluate potential antiviral role of the canonical Toll pathway components including MyD88, Tube, Pelle and Dorsal, shrimps with five groups were injected with each of the four components dsRNAs and the GFP dsRNA as control, respectively. Forty-eight hours later, each shrimp from the five groups was challenged with 10^5^ copies of WSSV particles and the gill tissues from 8 shrimps were sampled at 48 hours post infection to examine the virus copies by absolute qPCR. To explore the antiviral function of AMPs, a similar manipulation of RNAi plus WSSV challenge were performed with differences that gills were collected at more sampling times, orderly at the 24 h, 48 h, 72 h, 96 h and 120 h post infection.

To investigate the effects of Toll4 or Dorsal on the expression of AMPs *in vivo* after WSSV infection, AMPs expression in shrimps after receiving Toll4 dsRNA or Dorsal dsRNA plus WSSV challenge were analyzed. Hemocyte and/ or gill tissues from 9 shrimps were collected at 6 h post WSSV challenge, and the mRNA levels of fourteen AMPs were detected by qRT-PCR with specific primers (Supplement Table 1).

### Shrimp mortality or survival assay

Healthy shrimps were injected with gene specific dsRNAs including Toll4, ALF1, ALF2, ALF3, ALF4, LYZ1, LYZ2, LYZ3, LYZ4 or GFP dsRNA (as a control), and 48 h later were challenged with 10^5^ copies of WSSV particles in 50 μL PBS. Shrimps were kept in culture flasks for about 5 - 7 days following infection. The death of shrimp was recorded every 8 h and subjected to mortality or survival rate analysis.

### SDS-PAGE and western blotting

Hemocytes of normal shrimp and WSSV challenged shrimps were sampled with each sample collected and pooled from 5 shrimps. The nuclear and cytoplasmic fractions of hemocytes were extracted according to the protocol of NE-PER Nuclear and Cytoplasmic Extraction Reagents (Thermo, USA), while the total proteins were collected by RIPA lysis buffer. Samples were boiled for 5 min, separated on SDS-PAGE gels followed by transfer to polyvinylidene difluoride (PVDF) membranes. After blocking in 5% bovine serum albumin (BSA) in TBS with 0.1% Tween-20 (TBS-T) for 1 h, membranes were incubated with anti-Dorsal, anti-NF-κB p65 (phospho S276), anti-Cactus, anti-HSP90, anti-Histone H3 or anti-Actin for 16 - 18 h at 4 °C. After washing in TBS-T, membranes were incubated for 1 h at RT with horseradish peroxidase (HRP)-labeled Goat secondary antibody to Guinea pig IgG (H+L), Goat anti-Rabbit IgG (H+L)-HRP or Goat anti-Mouse IgG (H+L)-HRP. Both primary and secondary antibodies were incubated in TBS-T with 0.5% BSA. Membranes were developed with the enhanced chemiluminescent (ECL) blotting substrate (Thermo Scientific) and chemiluminescence was detected using the 5200 Chemiluminescence Imaging System (Tanon). For relative densitometry of Dorsal, Dorsal-P or Cactus, the immunoblotted band volume was normalized to the corresponding internal protein volume in the lane, using the ImageJ software 1.6.0 (National Institutes of Health, Bethesda, MD). Statistical analysis of densitometry data from three independent experiments was performed by using the Student’s t test.

### Pull-down assay

The *L. vannamei* AMPs ALF1 (accession No. AVP74301) and LYZ1 (ABD65298) without N-terminal signal peptide were cloned into pET-32a (+) plasmid (Merck Millipore, Germany) specific primers (Supplement Table 1), expressed in BL21 (DE3) *Escherichia coli* strain, and purified with Ni-NTA agarose (Qiagen, Germany) according to user’s manual. WSSV structural genes including VP19 (accession No. NP_477936.1) (77), VP24 (NP_477524.1) (78), VP26 (NP_477833.1) (79), VP28 (NP_477943.1) (79), wsv134 (NP_477656.1) (80) and wsv321 (NP_477843.1) (81) were cloned into pGEX-4T-1 plasmid (GE Healthcare, USA) with specific primers (Supplement Table 1), expressed in BL21 (DE3) *Escherichia coli* strain, and purified with Pierce™ GST agarose (Thermo Scientific) recommended by user’s operation. For His pull-down assay, purified His-tagged ALF1 or LYZ1 was incubated with Ni-NTA beads, to which purified WSSV structural protein was added and incubated at 4 °C for overnight with slight rotation. The mixture (beads and binding proteins) was washed three times with wash buffer (20 mM Imidazole, 50 mM Tris-HCl, pH 8.0), and then eluted in elution buffer (250 mM Imidazole, 50 mM Tris-HCl, pH 8.0). Elute was run in 10% SDS-PAGE, followed by coomassie staining and western blotting with anti-GST antibody to probe interacting proteins in the complex. For GST pull-down assay, purified GST-tagged WSSV structural protein and purified His-tagged ALF1 or LYZ1 were incubated with glutathione beads at 4 °C for overnight with slight rotation. The mixture was washed three times with PBS and the bound proteins were eluted in elution buffer (10 mM reduced glutathionem, 50 mM Tris-HCl, pH 8.0) and analyzed by SDS-PAGE as described above, followed by coomassie staining and western blotting with anti-His antibody.

### Immunocytochemical staning

Immunocytochemical staning was used to analysis shrimp Dorsal translocation in hemocyte recommended by a published method with a minor modification (46). In short, hemocytes were collected by centrifugation (1000 *g*, 5 min) at room temperature (RT) and deposited onto a glass slide, and then fixed immediately with 4% paraformaldehyde at RT for 5 min. The hemocytes on the glass slides was washed with PBS three times, followed by incubated with prepared Guinea pig anti-Dorsal antibody serum (1:2000 diluted in 5% BSA) overnight at 4 °C. The hemocytes were then washed with PBS and incubated with 5% BSA for 10 min; the Goat anti-Guinea pig IgG (H+L) Alexa Fluor 488 (Abcam, 1:5000 diluted in 5% BSA) was then added, and the samples were incubated for 1 h at RT in the dark. After being washed three times, the hemocytes were stained with Hochest (Sigma, 1 μg/ ml in PBS) for 10 min at RT and washed six times. Fluorescence was visualized on a confocal laser scanning microscope (Leica TCS-SP5, Germany). WCIF ImageJ software was used to analyze the colocalization of Dorsal and Hochest-stained nuclei in hemocytes according to a previously published method (46).

### Inhibitor injection

The QNZ (EVP4593) (S4902, Selleck) was reported to be a high-affinity partial antagonist of NF-κB (82, 83). Firstly, this NF-κB inhibitor with 0.5, 1.0 or 2 μg was injected into each shrimp (~8 g each) to explore the suppression effect on Dorsal. Then, the 2 μg NF-κB inhibitor for each shrimp was determined and used in the following treatments. DMSO injection was used as a control. The hemocytes of NF-κB inhibitor injected shrimp were sampled for protein and RNA extraction, as well as Dorsal translocation assay, at 6 h after WSSV challenge. The nuclear location of Dorsal was addressed by immunofluorescence staining as described earlier, the phosphorylation level of Dorsal was analyzed by western blotting with anti-NF-κB p65 (phospho S276) antibody, and the AMPs expressions were detected by qRT-PCR at 6 h after WSSV challenge. Besides, the gills were also collected for the AMPs expressions analysis by qRT-PCR.

### Statistical analysis

All data were presented as means ± SD. Student t test was used to calculate the comparisons between groups of numerical data. For mortality or survival rates, data were subjected to statistical analysis using GraphPad Prism software to generate the Kaplan–Meier plot (log-rank χ^2^ test).

---

**Figure S1.**
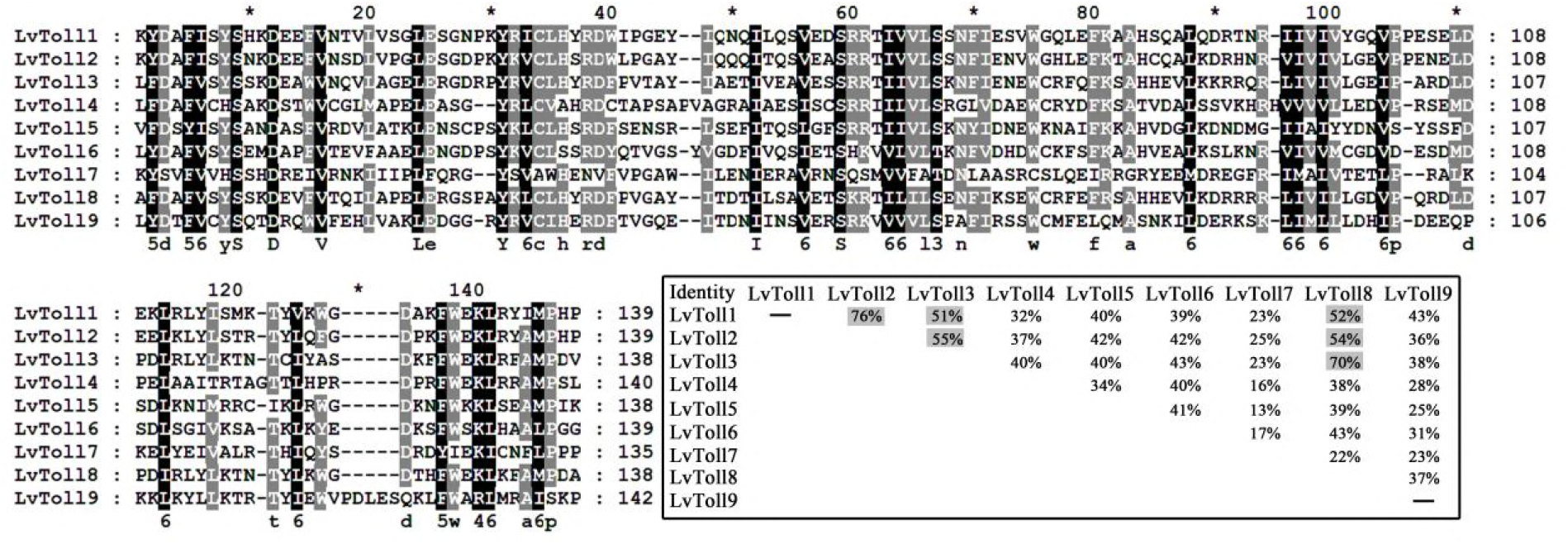
Multiple sequence alignment of the TIR domains of *L. vannamei* Toll1-9. The sequence identities among each other were calculated, and the values of greater than or equal to 50% were shaded.

## Supplement materials legends

**Supplement Data S1.** The cDNA sequences of nine *L. vannamei* Tolls (Toll1-9) including the 5’-untranslated region (UTR), 3’-UTR containing a poly (A) tail, and open reading frame (ORF) underlined.

**Supplement Data S2.** The cDNA sequences of fourteen *L. vannamei* AMPs including ALF1-4, LYZ1-4, PEN2-4 and CRU1-3. The open reading frames (ORFs) of these AMPs were underlined.

**Supplement Data S3.** The sequences of Tolls and Toll like receptors (TLRs) were used in the phylogenetic tree analysis, the TIR domains were underlined.

**Supplement Table 1.** Summary of primers in this study for dsRNA synthesis, Semi-quantitative reverse transcription PCR (Semi-qRT-PCR), quantitative reverse transcription PCR (qRT-PCR), absolute quantitative PCR and protein expression.

